# Expression of human ACE2 N-terminal domain, part of the receptor for SARS-CoV-2, in fusion with maltose binding protein, *E. coli* ribonuclease I and human RNase A

**DOI:** 10.1101/2021.01.31.429007

**Authors:** Shuang-yong Xu, Alexey Fomenkov, Tien-Hao Chen, Erbay Yigit

## Abstract

The SARS-CoV-2 viral genome contains a positive-strand single-stranded RNA of ~30 kb. Human ACE2 protein is the receptor for SARS-CoV-2 virus attachment and initiation of infection. We propose to use ribonucleases (RNases) as antiviral agents to destroy the viral genome *in vitro.* In the virions the RNA is protected by viral capsid proteins, membrane proteins and nucleocapsid proteins. To overcome this protection we set out to construct RNase fusion with human ACE2 receptor N-terminal domain (ACE2NTD). We constructed six proteins expressed in *E. coli* cells: 1) MBP-ACE2NTD, 2) ACE2NTD-GFP, 3) RNase I (6xHis), 4) RNase III (6xHis), 5) RNase I-ACE2NTD (6xHis), and 6) human RNase A-ACE2NTD150 (6xHis). We evaluated fusion expression in different *E. coli* strains, partially purified MBP-ACE2NTD protein from the soluble fraction of bacterial cell lysate, and refolded MBP-ACE2NTD protein from inclusion body. The engineered RNase I-ACE2NTD (6xHis) and hRNase A-ACE2NTD (6xHis) fusions are active in cleaving COVID-19 RNA *in vitro.* The recombinant RNase I (6xHis) and RNase III (6xHis) are active in cleaving RNA and dsRNA in test tube. This study provides a proof-of-concept for construction of fusion protein between human cell receptor and nuclease that may be used to degrade viral nucleic acids in our environment.

**Graphical Abstract:** Cartoon illustration part of this work (Human ACE2 N-terminal domain tethered to RNase A and RNA degradation by the fusion enzyme).

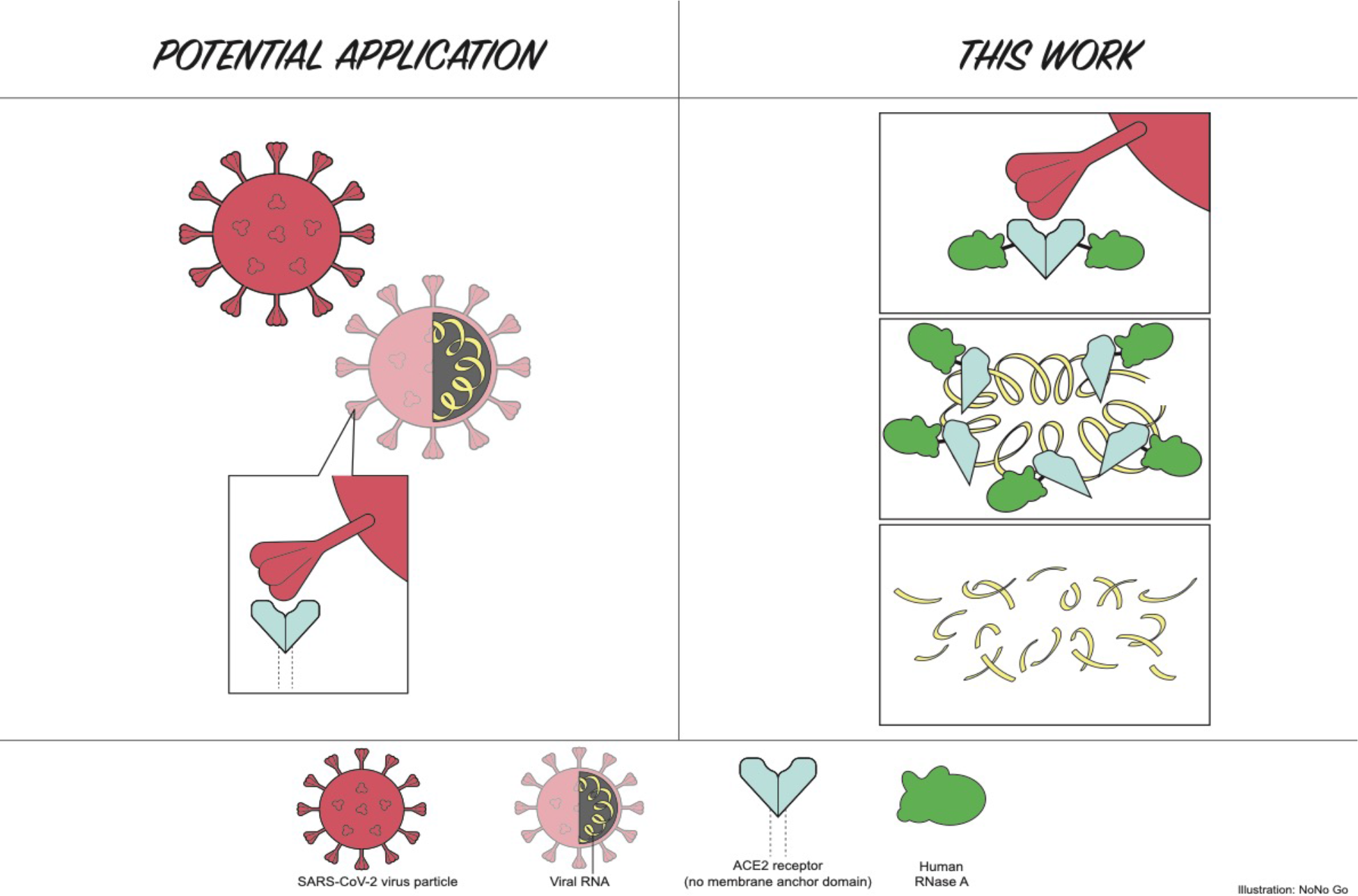

## Introduction

Human angiotensin-converting enzyme 2 (ACE2 or ACE-2) is a carboxypeptidase which converts angiotensin I to angiotensin 1-9, and it catalyzes the degradation of angiotensin II to angiotensin 1-7 (reviewed in [1; 2]), which attenuates inflammation and oxidative stress caused by angiotensin II. ACE2 is a glycoprotein that serves as host cell receptor for the severe acute respiratory syndrome coronavirus (SARS-CoV), human coronavirus NL63/HCoV-NL63, and SARS-CoV-2 [3; 4]. The SARS-CoV-2 is currently causing pandemics and COVID-19 respiratory disease around the globe since its initial outbreak in Wuhan, Hubei Province of China in December 2019 [5]. The first genome sequence was deposited in GenBank/NCBI on Jan. 13, 2020 (Wuhan seafood market pneumonia virus isolate Wuhan-Hu-1, GenBank accession number NC_045512). The mortality rate among the elderly (>65) with underlying conditions is high, while the overall fatality rate from positively diagnosed patients in the world stands at ~6.26 % as of April 14, 2020 (120,863 death among over 1.9 million infected when this work was half-finished) [6]. For comparison, the global mortality associated with seasonal influenza epidemics from 2002-2011 (no data in 2009) from World Health Organization was estimated at 0.00058%, or approximately 6 in 10,000 infected individuals in all age groups [7]. Thus, COVID-19 is approximately 10-100 times more lethal in causing human death than seasonal flu, although the mortality rate varies from country to country depending on the medical infrastructure. In majority of COVID-19 positive cases (estimated at ~70% to 80%), the infected individuals show symptoms with fevers and chill, severe muscle pain, cough, symptoms of upper respiratory tract infection and will likely recover after 2-3 weeks. However, the other 20%-30% patients require hospitalization and roughly 5% patients become critically ill and require intensive care and some would eventually develop severe pneumonia (acute respiratory distress syndrome), cardiovascular damage, acute kidney injury, and catastrophic organ failure, possibly owing to acute cytokine storm of massive host immunological reactions [8; 9; 10]. Autoantibodies against immune cells and interferons are also associated with severe COVID-19 disease. The percentage of asymptomatic positive carriers is unknown due to the lack of extensive antibody testing in the general population. The soluble recombinant human ACE2 protein has been shown to drastically reduce virus titer in cell cultures by binding to SARS-CoV-2 virus and preventing reinfection [11].

The SARS viral spike (S) protein forms a trimer and it’s erected active form mediates virus entry into host cells by first binding to the ACE2 receptor through the receptor-binding domain (RBD) in the N-terminal S1 domain (the trimeric Spike protein also binds cellular heparin). Following protease cleavage, fusion of the viral and host membranes through the S2 domain ensued by endocytosis and subsequent release of viral RNA for translation of viral proteins, post-translational processing, viral RNA replication and modification, and virus particle maturation in Golgi apparatus [12; 13; 14; 15]. The life cycle of SARS-CoV-2 is probably very similar to that of SARS-CoV [4; 16]. The receptor binding domain of the S protein (S-RBD) serves as a molecular target for developing virus attachment inhibitors, therapeutic antibodies, and preventive vaccines [17]. The structures of ACE2 receptor with bound S1 domain or fulllength S protein of SARS-CoV-2 have been solved recently at high resolution by Cryo-EM [18; 19]. Recently, COVID-19 virus variants have been isolated in UK, South Africa, Brazil, and US that contain multiple amino acid (aa) mutations in the S protein (e.g. K417N, E484K, N501Y aa substitutions) that may have increased the binding affinity to human ACE2 receptor and enhanced transmission rate [20]. Another host protein, the transmembrane protease, serine 2 (TMPRSS2) is believed to cleave the SARS-CoV-2 spike protein to facilitate cellular attachment and membrane fusion [4; 21]. While cellular proteases furin and cathepsin B/L proteases can also cleave S protein and active the virus for reinfection of other cells. The full-length TMPRSS2 protein is 492-aa long, that can be cleaved into two polypeptides by autocatalysis: N-terminal non-catalytic domain (aa 1-255), and C-terminal catalytic domain (aa 256-492). The Serine protease facilitates the phylogenetically related SARS-CoV virus-cell membrane fusions by proteolytically cleaving and processing the viral spike glycoproteins [22]. Over-expression of TMPRSS2 in human VeroE6 cells resulted in a cell line VeroE6/TMPRSS2 highly susceptible to SARS-CoV-2 infection and potentially useful for generating high virus titers [23].

Current strategies to reduce viral transmission among individuals are social distancing (curfew, self-quarantine and/or mandatory shelter in place) to reduce community transmission, chemical treatment and sanitization of public places that may harbor the viral particles, creation of physical barriers (wearing masks, facial shield, protective goggle and gown for health care workers). The SARS-CoV particles on smooth surfaces retained their viability for over 5 days at 22-25°C and relative humidity of 40-50%, however, virus viability was rapidly lost at higher temperatures (>38°C) and higher relative humidity [24]. The survival of infectious SARS-CoV-2 virus particle in air droplets and solid surfaces is estimated in a few hours to days [25]. Virus particles in the respiratory air droplets (aerosol) from asymptomatic carriers and patients are suspected as transmission route, which resulted in high transmission rate to health care workers in hospitals and other high density social gathering. The degradation and loss of infectivity of the virus particles relies on natural decays of the infectious particles possibly by many environmental factors such as temperature, humidity, pH, protease and RNase and other natural microbial enzymes present under environmental settings. Three vaccines against SARS-CoV-2 Spike (S) protein have been developed by pharmaceutical companies and are being applied to front line medical professionals, senior citizens, essential workers and later to the general population [26; 27; 28; 29; 30; 31; 32]. Phase 1 clinical trial for a single dose intranasal vaccine was initiated in Q4 2020 by Altimmune Inc. and University of Alabama/Birmingham.

Clinical trials have been carried out for treatment of COVID-19 with the nucleotide analog RNA-dependent RNA Polymerase (RdRP) inhibitor Remdesivir^®^, which has been approved by the US Food and Drug Administration to treat COVID-19 patients. Other antiviral drugs include protease inhibitors that inhibit the viral protease (main protease, Mpro) required for the processing of virus polyprotein into functional enzymes (RdRP, RNA helicase, and ribonuclease) for viral genome replication. Other host factors involved in viral replication and maturations have been identified and are being targeted for therapeutic intervention [33]. The overall economic fallout from the SARS-CoV-2 pandemics in the global economy is difficult to quantitate, but the pandemics had negatively impacted on economic activities and GDP, leading to a global recession in 2020.

The *E. coli* MBP protein expression system utilizes a strong promoter for transcription and a strong ribosome binding site for protein synthesis and the production of target protein is inducible by addition of IPTG (www.neb.com)[34]. Some MBP fusion proteins can be expressed at high level, up to 100 mg per liter (L) of induced cells. In addition, the target protein in fusion with MBP can be exported into bacterial periplasm to avoid toxicity since over-expression of certain enzymes in cytoplasm is toxic to the host. MBP signal peptide directed fusion protein to periplasm (removal of MBP signal peptide). In some cases, MBP fusion with the target protein is active and there is no need to cleave off the MBP tag for enzymatic activity (e.g. MBP-PNGase F (NEB Catalog 2019/20), RNase I_f_ (MBP-RNase I fusion), MBP-*Geobacillus* intron-encoded reverse transcriptase (Xu, unpublished). The MBP tag can also serve as a chaperon to enhance the fusion protein solubility [34; 35]. The *E. coli* ribonuclease I (RNase I, periplasmic isoform) belongs to the RNase T2/S-RNase group of endoribonucleases and it had been over-expressed and purified [36]. Bovine RNase A and *E. coli* RNase I, commercially available in research grade for many years, are widely used to eliminate RNA in preparation of plasmid and genomic DNA for molecular biology applications [37]. *E. coli* RNase III is an endoribonuclease cleaving double-stranded (ds) duplex RNA that is involved in transcript degradation and accelerated RNA decay and it requires divalent cations for activity [37]. RNase III and its variants can generate small dsRNA (21-23 bp) in Mn^2+^ buffer to be used in small RNA-medicated gene transcription regulation [38]. The human ribonuclease A (hRNase A) has eight different secreted variants (isoforms) that have antiviral/antibacterial/antifungal activities, and cytotoxicity against intracellular parasites. hRNase A (pancreatic) encoded by the human *rnase1* gene is an endorinuclease cleaving the 3’ side of pyrimidine nucleotides of single-stranded and doublestranded RNA [39; 40]. The hRNase A precursor contains 156 aa residues with a signal peptide of 28 residues (128 aa mature protein, Uniprot ID-P07998). The potential of hRNase A as an anti-Covid-19 reagent has not been explored to our knowledge.

The SARS-CoV-2 viral genome is a positive-strand ssRNA of ~30 kb possibly encoding at least 29 viral proteins [41]. In this study, we propose to use ribonucleases (RNases) as an antiviral agent to destroy the viral genome. Our goal is to construct RNase I fusion with human ACE2 receptor protein and to investigate strategies to trigger the destabilization of the viral particles or partially release the RNA genome to become more accessible to RNase degradation. To achieve this goal, we expressed six proteins in *E. coli* cells: 1) MBP-ACE2NTD, 2) ACE2 NTD-GFP, 3) RNase I (6xHis), 4) RNase III (6xHis), 5) RNase I-ACE2NTD (6xHis), 6) hRNase A-ACE2NTD. We examined expression conditions and partially purified the fusion protein MBP-ACE2NTD. We optimized expression conditions and strains for RNase I (6xHis) and RNase III (6xHis). We also constructed and partially purified the fusion enzyme RNase I-ACE2NTD (6xHis) which is active in cleaving RNA in the absence of divalent cations. We also demonstrated that hRNase A-ACE2NTD (6xHis) fusion is active in digestion of Covid-19 RNA fragment and other RNA substrates. We fused ACE2-NTD to GFP that resulted in reduced green fluorescence signal. This construct may be useful to isolate more soluble variants of ACE2NTD by screening mutant libraries. This work serves as a proof of concept for the construction of artificial ribonuclease fused to human ACE2 receptor NTD. The availability of inexpensive recombinant ACE2NTD protein may facilitate the screening of small molecules binding to ACE2 and block SARS virus attachment and entry.

## Materials and Methods

### E. coli strains and recombinant DNA method

*E. coli* strains for fusion protein expression are NEB Express (B strain), NEB Turbo (K strain), NEB SHuffle^®^ T7 (K and B strains), Nico (λDE3) derived from BL21. The periplasm expression vector pMAL-p5x (Amp^R^) was used for the construction of ACE2NTD fusion. Synthetic gene blocks (gblock) with optimized *E. coli* codons were purchased from Integrated DNA Technology (IDT) and cloned into the expression vector (NdeI and BamHI digested) by NEB HiFi assembly enzyme mix (E2621S, NEB, MA) according to the manufacturer’s instruction. Assembled vector with potential insert was transferred into *E. coli* competent cells by transformation. Individual Amp^R^ transformants were picked and cultured in 2 ml of LB + Amp and shaken at 37°C for 5-6 h. 1.5 ml of cells were spun down and kept in −20°C for plasmid mini-preparation until needed. 10 ml of LB + Amp were added to the remaining cells and cultured for ~2-3 h to reach late log phase. IPTG at 0.3 mM final concentration was added for fusion protein production overnight (at 16°C to 18°C). Plasmid DNA was prepared by a Monarch^®^ mini-preparation kit (NEB). The inserts were sequenced by two primers (forward primer S1273 from the C-terminal coding region of MBP, and a reverse primer (R2913) downstream of the BamHI site of pMAL-p5x) with a BigDye^®^ terminator V3.1 cycle sequencing kit (ABI/Thermo-Fisher). Amylose resin (E8021S, NEB) and amylose column chromatography was carried out according to the manufacturer’s protocol. Protein refolding kit was purchased from Novagen. Proteins in pellet (inclusion body) after centrifugation (10,000 xg) was resuspended in a solubilization buffer (10 mM CAPS, pH 11.0, 0.3% N-lauroylsarcosine) and pelleted in a microcentrifuge (4°C). After washing the pellet with wash buffer twice (20 mM Tris-HCl, pH 7.5, 10 mM EDTA, 1% Triton X-100) the pellet was solubilized and dialyzed against 2 L of dialysis buffer (20 mM Tris-HCl, pH 8.5, 1 mM DTT) for 4 h to overnight at 4°C with exchange of the dialysis buffer twice. A small fraction of the refolded MBP-ACE2NTD fusion protein was further purified by amylose magnetic beads (NEB) according to the protocol provided by the manufacturer.

### Description of the amino acid (aa) sequence of MBP-ACE2NTD fusion

The full-length ACE2 protein has 805 aa residues (GenBank sequence ID: BAB40370.1). The first 17 aa residues form the signal peptide not required for activity; the extracellular domain is consisted of aa residues 18-740 [42]. Therefore, only the coding sequence for aa residues 12 to 444 is synthesized in two gene blocks (IDT) (the first 11 aa residues were deleted, #12 Val codon was changed to ATG codon for cloning purpose. See the modified protein sequence below). The S2 domain involved in membrane fusion is not included here.

Maltose binding protein (MBP)-ACE2NTD (NdeI)

MSSSSWLLLS LMAVTAAQST IEEQAKTFLD <u>KFNHEAEDLF Y</u>QSSLASWNY

NTNITEENVQ NMNNAGDKWS A<u>FLK</u>EQSTLA QMYPLQEIQN LTVKLQLQAL

QQNGSSVLSE DKSKRLNTIL NTMSTIYSTG KVCNPDNPQE CLLLEPGLNE

IMANSLDYNE RLWAWESWRS EVGKQLRPLY EEYVVLKNEM ARANHYEDYG

DYWRGDYEVN GVDGYDYSRG QLIEDVEHTF EEIKPLYEHL HAYVRAKLMN

AYPSYISPIG CLPAHLLGDM WGRFWTNLYS LTVPFGQKPN IDVTDAMVDQ

AWDAQRIFKE AEKFFVSVGL PNMTQGFWEN SMLTDPGNVQ KAVCHPTAWD

LG<u>*K*GDFR</u>ILM CTKVTMDDFL TAHHEMGHIQ YDMAYAAQPF LLRNGANEGF

HEAVGEIMSL SAATPKHLKS IGLLSPDFQE DNETEINFLL KQAL(BamHI)

(The aa residues for Spike protein binding are underlined)

The codon-optimized gblocks were assembled into pMAL-p5x (p for periplasmic expression), which encodes MBP-ACE2NTD fusion with predicted molecular mass (MW) of ~94.0 kDa (mature protein had 26-aa signal peptide removed from MBP N-terminus, 44.5 + 49.5 = 94.0 kDa). The first six aa residues (MAVTAA) serve as a short linker between MBP and ACE2NTD. SARS-CoV-2 His-tagged Spike (S) protein and the receptor binding domain (RBD) protein of S were purchased from Sino Biological (Beijing/China and Wayne/USA).

### Construction of ACE2NTD-GFP fusion

The coding sequence for ACE2NTD was amplified by PCR and inserted into pDasherGFP (NdeI cut, pBR322 backbone, Amp^R^)(source reference: atum.bio). GFP expression was under the control of T5 promoter and is inducible by addition of IPTG. GFP expressing colonies, GFP and ACE2NTD-GFP protein bands were visualized under long UV light or by a fluorescence imager (Typhoon, GE Lifesciences) at 520 nM (Cy2 channel, 515-535 nm emission).

### Cloning of E. coli RNase I coding sequence into pET21b with a C-terminal 6xHis tag and construction of RNase I-ACE2NTD fusion

The RNase I gene (rna) was amplified from *E. coli* genomic DNA (*E. coli* strain H709c) and assembled into T7 expression vector pET21b (NdeI and XhoI digested), which produced a C-terminal 6xHis tag. The protein expression level was evaluated in four T7 expression strains: T7 Express (C2566, NEB), T7 Express with LysY and *lacI^q^* (C3013, NEB), T7 SHuffle (C3026, *E. coli* K strain, NEB) and Nico (λDE3) with reduced histidine-rich background proteins (NEB). The RNase I precursor contains a signal peptide (amino acid residues 1-23) for export to periplasm. To create RNase I-ACE2NTD fusion, a PCR fragment amplified from ACE2NTD gene block (IDT) was assembled into the XhoI site of pET21b-RNase I, which generated the coding sequence for the fusion RNase I-ACE2NTD (6xHis). The entire gene was sequenced using T7 universal primer(S1248), T7 terminator primer (reverse) (S1271), and internal ACE2 primers. The expression of RNase I-ACE2NTD (6xHis) fusion was evaluated in five T7 expression strains: C2566, C3013, C3026, Nico (λDE3), and BL21 (λDE3). RNase I (6xHis) was partially purified by chromatography through Ni-NTA column (Ni-NTA agarose fast-flow, ThermoFisher). The protein was concentrated by low-speed centrifugation in an Amicon concentrator (10 kDa cut-off) and resuspended in a storage buffer SB (50 mM Tris-HCl, pH 7.5, 200 mM NaCl, 10 mM DTT, 50% glycerol).

### Cloning of E. coli RNase III gene (rnc) into pET21b with a C-terminal 6xHis tag

The RNase III gene (rnc) was amplified from *E. coli* genomic DNA (*E. coli* strain H709c) and assembled into T7 expression vector pET21b (NdeI and XhoI digested), which produced a C-terminal 6xHis tag. The protein expression level was evaluated in three T7 expression strains: T7 Express with LysY and *lacE* (C3013), T7 SHuffle (C3026, *E. coli* K strain) and Nico (λDE3). RNase III (6xHis) was partially purified by chromatography through Ni-NTA column (Ni-NTA agarose fast-flow, ThermoFisher). The protein was concentrated by low-speed centrifugation in a concentrator and resuspended in a storage buffer SB. RNase III activity was assayed using a 40mer duplex RNA and dsRNA ladder (NEB).

### Cloning of E. coli RNase I (no native signal peptide coding sequence) gene into pMAL-p5x

A PCR fragment encoding *E. coli* RNase I (GenBank Sequence ID: AAB40811.1) was inserted into pMAL-p5x to generate the following construct (the 23-aa signal sequence of RNase I was deleted). The molecular weight of the fusion protein MBP-RNase I is predicted to be ~71.7 kDa (44.5 + 27.2 kDa).

(P_tac_)-MBP-(NdeI)-

(MKAFWRNAALLAVSLLPFSSANA) LALQAKQYGDFDRYVLALSWQTGFCQSQHDRNRNERD

ECRLQTETTNKADFLTVHGLWPGLPKSVAARGVDERRWMRFGCATRPIPNLPEARASRMC

SSPETGLSLETAAKLSEVMPGAGGRSCLERYEYAKHGACFGFDPDAYFGTMVRLNQEIKE

SEAGKFLADNYGKTVSRRDFDAAFAKSWGKENVKAVKLTCQGNPAYLTEIQISIKADAIN

APLSANSFLPQPHPGNCGKTFVIDKAGY (BamHI)

### Cloning of human RNase A gene (human rnase1) and construction of hRNaseA-ACE2NTD150 fusion

Two versions of fusion were constructed: fusion 1 contains the *E. coli* MBP signal peptide (aa residues 1-26, this signal peptide directed the fusion protein to be exported to the periplasmic space), and fusion 2 without the MBP signal peptide. The fusion is consisted of peptides: *E. coli* MBP signal peptide (MBP residues 1-26) + hRNase A (hRNase A residues 29-156) + ACE2NTD (ACE2 N-terminal residues 18-150, referred to as ACE2NTD150) (note: the aa residue numbers refer to the native protein amino acid number).

MKIKTGARIL ALSALTTMMF SASALA (*E. coli* MBP signal peptide)

KESRAKKFQRQHMDSDSSPSSSSTYCNQMMRRRNMTQGRCKPVNTFVHEPLVDVQNVCFQEKVTCKNGQGNCYKSNSSMHITDCRLTNGSRYPNCAYRTSPKERHIIVACEG SPYV PVHFDASVEDST

(hRNase A)

<u>GSAGSA</u> (artificial linker aa)

QSTIEEQAKTFLDKFNHEAEDLFYQSSLASWNYNTNITEENVQNMNNAGDKWSAFLKE

QSTLAQMYPLQEIQNLTVKLQLQALQQNGSSVLSEDKSKRLNTILNTMSTIY

STGKVCNPDNPQECLLLEPGHHHHHH (ACE2NTD150-6xHis).

The predicted molecular mass of hRNaseA-ACE2 (6xHis) with and without MBP signal peptide is 33.3 kDa and 30.7 kDa, respectively. The fusion gene blocks were purchased from IDT and assembled into T7 expression vector pET21b (NdeI and XhoI cut), which resulted in a C-terminal 6xHis tag. The protein expression level was evaluated in three T7 expression strains: T7 Express with LysY and *lacI^q^* (C3013), T7 SHuffle (C3029, *E. coli* B strain) and Nico (λDE3, C2529H). Soluble hRNaseA-ACE2 (6xHis) fusion without signal peptide was purified from 1 L of IPTG-induced cells (cells induced at 16°C overnight) by chromatography through a Ni-NTA agarose column or from 3 ml of cell lysate (30 ml of IPTG-induced cells) by binding to Ni magnetic beads.

### RNase I and RNase III activity assays

RNase III assay: 1 μg double-stranded (ds) RNA substrates were digested with either ShortCut^®^ RNase III (NEB) or RNase III (6xHis) for 15 min at 37°C in total volume of 15 μL 1x reaction buffer (50 mM Tris-HCl, 1 mM DTT, 50 mM NaCl, pH 7.5) supplemented with 20 mM MnCl2, and stopped by addition EDTA (50 mM). Cleavage products were analyzed by gel electrophoresis on a 2% agarose gel.

ssRNA, dsRNA ladders and a 5’ Fluorescein-labeled RNA (300 nt) were provided by NEB and used as substrates for RNase I and RNase III activity assays. Cleavage products were analyzed by gel electrophoresis on PAGE-Urea gel or agarose gel. 5’ FAM-labeled COVID-19 ssRNA (60 mer) was purchased from IDT. RNase I assay: 16 nM of the RNA substrate was digested by purified MBP-RNase I, RNase I (6xHis) or refolded RNase I-ACE2NTD (6xHis) in a high salt buffer (NEB buffer 3: 100 mM NaCl, 50 mM Tris-HCl, pH 7.9 at 25°C, 10 mM MgCl_2_, 1 mM DTT) or a high salt buffer without divalent cations (100 mM NaCl, 50 mM Tris-HCl, pH 7.5, 1 mM DTT) as RNase I is active in the absence of divalent cations or in the presence of EDTA [37]. Cleavage reactions were quenched by eliminating enzymes with the treatment with Proteinase K (1.6 U, NEB) and the products were analyzed by capillary gel electrophoresis (CE assay). Substrate and cleavage product peaks were analyzed by PeakScan software (ABI/ThermoFisher).

### Protein pull-down assays (protein complex pull-down by Ni magnetic beads)

#### Ni-magnetic beads pull down of His-tagged S protein in complex with MBP-ACE2NTD

Ni-magnetic beads (100 μl) was washed three times with Ni binding buffer (50 mM NaH2PO4, 0.3 M NaCl, 20 mM imidazole). His-tagged spike protein (2 μg, purchased from Sino Biological) was added to the beads pre-suspended in 200 μl Ni binding buffer and incubated at 4°C in a roller for 30 min. Refolded MBP-ACE2NTD protein (10 μg) or equal volume of storage buffer was added to the prebound Ni-beads/S protein and co-incubation continued at 4°C for 1 h. The Ni magnetic beads were washed three times with Ni binding buffer (1 ml each time) and the protein was eluted by addition of 50 μl of Ni elution buffer (50 mM NaH2PO4, 0.3 M NaCl, 0.25 M imidazole). Approximately 17 μl of the eluted protein was loaded onto 10-20% SDS-PAG gel for analysis.

#### Amylose magnetic beads pull down of His-tagged S protein and RBD protein

NEB’s protocol for MBP protein binding to the amylose magnetic beads was used with minor modification. Amylose magnetic beads (400 μl) in an Eppendorf tube were saturated (blocked) in 1 ml of BSA (1x) in MBP binding buffer (200 mM NaCl, 20 mM Tris-HCl, pH 7.4, 1 mM EDTA, 1 mM DTT) on a rotator at 4°C overnight. The beads were then washed 3 times (15 min each) at 4°C with MBP binding buffer. The magnetic beads were resuspended in 200 μl of MBP binding buffer. His-tagged SARS-CoV-2 Spike protein (5 μl) or His-tagged RBD protein (5 μl at 1.75 μg/μl, purchased from Sino Biological) were first diluted in 45 μl of MBP binding buffer. MBP-ACE2 fusion protein (25 μl at 0.98 μg/μl) were mixed with diluted Spike protein or RBD protein and 100 μl of BSA-saturated magnetic beads. Protein binding (complex formation) was carried out on a rotator at 4°C overnight. The beads with bound proteins were washed 3 times with 1 ml of MBP binding buffer at 4°C for 15 min. Bound proteins were eluted with addition of 50 μl of MBP elution buffer (MBP binding buffer plus 10 mM maltose) and the eluted proteins were analyzed by Western blot using anti-His Ab or anti-ACE2 Ab.

### Western blot using anti-6xHis antibody (Ab) and mouse monoclonal anti-ACE2 Ab

Mouse monoclonal anti-ACE2 Ab and mouse anti-6xHis Ab were purchased from Cell Signaling Technologies (CST, MA) and EMC Millipore company, respectively. Protein expression, purification and interaction between protein partners were monitored by Western blot analysis according to the manufacturer’s protocol. Briefly, cell extract or purified proteins (6xHis tag) were subjected to 10-20% SDS PAGE and proteins were transferred onto Protran BA85 nitrocellulose (Whatman) in a 0.5x transfer buffer. The transferred proteins were stained with Ponceau S solution (Sigma-Aldrich). The nitrocellulose membrane was soaked (blocked) in 3% milk (dry milk from Bio-Rad) solution in 1x PBS following by incubation with appropriate primary antibody solution (1:1000 dilution) for 1 h at room temperature. The excess primary antibodies were washed off with 1x PBS containing 0.1% Tween 20 and 0.1% Triton X100 three times for 15 min, followed by incubation with secondary anti-mouse IgG HPR conjugated antibodies (1:1000) (CST) and secondary Ab, washed three times with 1x PBS, 0.1% Tween20 and 0.1% Triton X100. Signal detection was performed by treatment with LumiGlo reagent (CST) and visualized by fluorescent imaging.

## Results and Discussion

### Expression and purification of MBP-ACE2NTD fusion protein

A schematic diagram of the fusion consisting of an N-terminal MBP and a C-terminal ACE2NTD as well as the cloning sites is shown in **Fig. 1A**. Total cellular proteins from sonicated cell lysate of IPTG-induced cells (NEB Express, 16°C overnight) were analyzed by SDS-PAGE (**Fig. 1B)**. Nine out of 11 clones produced a strong protein band of ~94 kDa, close to the predicted size. Clone #8 contains the correct sequence and was chosen for further analysis. After 15 min centrifugation of the total lysate in a microcentrifuge, however, most of the MBP-ACE2NTD fusion was found in the pellet (inclusion body) (data not shown). To improve protein folding and solubility, NEB SHuffle *E. coli* host was tested for protein induction. This strain was engineered to promote disulfide bond formation in the cytoplasm and is deficient in proteases Lon and OmpT. 1 L cell culture was induced with IPTG at 16°C overnight and the supernatant of cell lysates (the soluble fraction) was loaded onto a 15 ml amylose column. **Fig. 1C** shows the partially purified fusion protein using an amylose column (~6 mg protein purified from 1 liter LB of IPTG-induced cells including a few contaminating proteins). The ~40 kDa protein is likely to be the host MBP without the signal peptide. To remove this major contaminating protein, an MBP-deficient SHuffle strain will be required. Some fusion protein expressed in NEB SHuffle strain was found in the pellet (**Fig. 1C**, lane 4). The protein pellet from NEB Express cells was solubilized. After solubilization and refolding, the fusion protein was partially purified (**Fig. 1C**, lane 5). The protein yield from solubilized pellet (NEB Express) was approximately 50 mg/L of IPTG-induced cells. The refolded protein can be further purified by incubation with amylose magnetic beads (**Fig. 1D**). It was concluded that most of the fusion protein made in NEB Express was found in the pellet (inclusion body). 1 L of IPTG-induced NEB SHuffle cells produced ~6 mg of soluble protein and ~50 mg of insoluble protein in the pellet. The production of MBP-ACE2NTD protein from *E. coli* cells may have a cost advantage over ACE2 protein produced from human cell cultures. But the ACE2NTD from bacterial source lacks the sugar modifications compared to the human version.

**Fig. 1.**
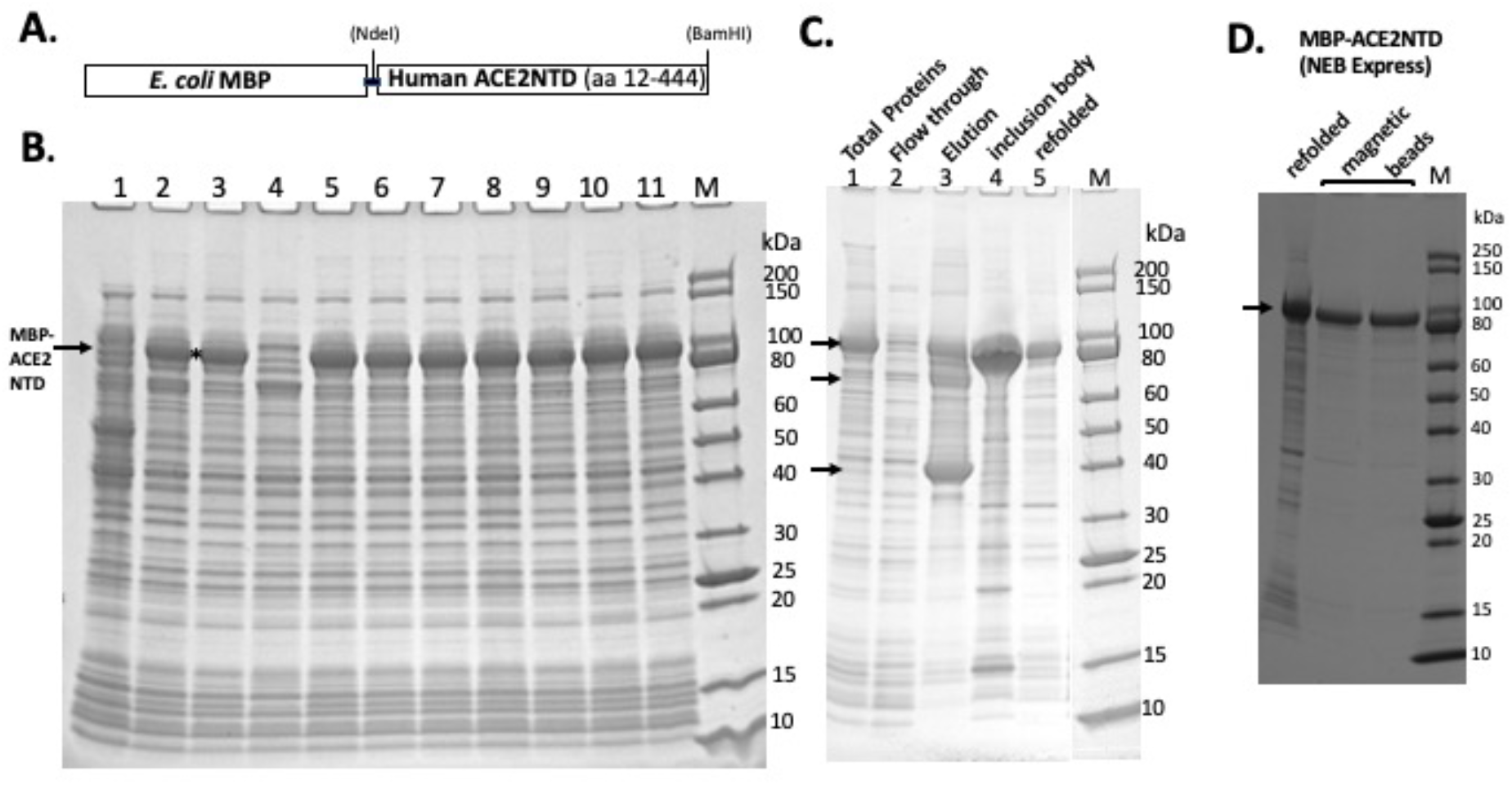
SDS-PAGE analysis of MBP-ACE2NTD fusion. **A.** A schematic diagram of the fusion. **B.** Total cellular proteins from IPTG-induced cells (NEB Express) containing the MBP-ACE2NTD fusion. 9 out of 11 clones tested here contained the fusion (except #1 and #4). Clone #8 was chosen for further study. **C.** MBP-ACE2NTD fusion expressed from IPTG induced cells (T7 Shuffle, lanes 1-4). Lane 1, total cellular protein; lane 2, flow-through from an amylose column; lane 3, eluted MBP-ACE2NTD protein from an amylose column; lane 4, MBP-ACE2NTD protein found in the pellet (inclusion body); lane 5, refolded MBP-ACE2NTD fusion protein from the pellet (NEB Express) shown in **B**; M, protein size marker (NEB). Arrows indicate the MBP-ACE2NTD protein, a truncated protein, and host MBP. **D.** Purified MBP-ACE2NTD protein by amylose magnetic beads. Lane 1, refolded fusion; lanes 2 and 3, protein eluted in a Tris-HCl buffer (20 mM, pH 7.5) and sodium phosphate buffer (0.1 M, pH 8.0) with 10 mM maltose.

### MBP-ACE2NTD binding to Spike (S) protein and receptor binding domain (RBD) of S protein

One potential application of MBP-ACE2NTD protein is to serve as a decoy to bind SARS-CoV-2. We examined MBP-ACE2NTD binding to S and RBD protein *in vitro* by protein pull-down assay. The His-tagged S protein is expected to bind to the Ni magnetic beads. The MBP-ACE2NTD protein was incubated with the S protein pre-bound to the Ni magnetic beads. After extensive washing with high salt buffer, the S protein and potential bound partner were eluted with an elution buffer. In a control experiment, approximately 20% of the S protein can be recovered from the Ni magnetic beads (**Suppl. Fig. S1 A**. compared lanes 1-input and lane 2-recovered). When the S protein was incubated with MBP-ACE2NTD protein, only a very weak band of the binding partner was recovered, indicating poor binding of S protein to MBP-ACE2NTD *in vitro* under the high salt condition (0.3 M NaCl). In a reversed pull-down assay, the MBP-ACE2NTD bound to the amylose magnetic beads was incubated with S or RBD proteins. After extensive washing, the proteins were eluted with a maltose buffer. Only a weak band of the S protein was detected by anti-6xHis Ab in the Western blot (**Suppl. Fig. S1 B**). The RBD protein binds to the binding partner modestly well, as detected by anti-6xHis Ab in the Western blot. In the control experiment, MBP-ACE2NTD was detected by monoclonal anti-ACE2 Ab. The protein binding and pull-down results indicate that the SARS-Cov-2 S protein binds poorly to the MBP-ACE2NTD protein, and the RBD protein binds reasonably well to the fusion in vitro. This result is consistent with published data that the SARS-CoV-2 S protein, unlike SARS-CoV-1 S protein, is a poor binder to the human ACE2 receptor *in vitro* [18; 19; 42]. It remains to be seen whether some recent SARS-CoV-2 variants (e.g. N501Y, P681H, and ΔH69/ΔV70 variants, P.1 variant with K417T/E484K/N501Y amino acid changes in S protein) with higher transmission rate can bind to the ACE2 receptor protein more tightly *in vitro* [20] (P.1 variant reference: https://virological.org/t/sars-cov-2-reinfection-by-the-new-variant-of-concern-voc-p-1-in-amazonas-brazil/596).

### ACE2NTD-GFP fusion

Since ACE2NTD fusion to MBP displayed reduced solubility, it is anticipated that fusion of ACE2NTD to GFP may reduce green fluorescence. ACE2NTD-GFP fusion was constructed by insertion of ACE2NTD coding sequence into the GFP expression vector, creating N-terminal ACE2NTD and C-terminal GFP. GFP-expressing *E. coli* cells (IPTG-induced) showed strong green fluorescence and formed green colonies under long UV light or at 520 nm light emission (**Suppl. Fig. S2 A**). The ACE2NTD-GFP expression cells (IPTG-induced NEB Express and T7 Express), however, did not show green fluorescence, presumably the ACE2NTD-GFP fusion negatively affected the protein folding and/or expression inside cells. IPTG induced expression of ACE2NTD-GFP fusion showed a weak green fluorescence in NEB SHuffle cells, suggesting improvement of folding of the fusion protein. To further confirm this, the total proteins from IPTG induced cells were analyzed by SDS-PAGE and protein bands were visualized on a fluorescence imager. GFP showed a strong fluorescence signal, but the ACE2NTD-GFP fusion protein showed a weak signal in NEB SHuffle K and B strains (**Suppl. Fig. S2 B**) (the fluorescence signal presumably resulted from protein refolding after washing three times in Milli-Q water). It was concluded that ACE2NTD likely negatively affected the folding and/or expression of GFP when they were fused together. This fusion system may be used to isolate ACE2NTD variants with improved folding and stronger fluorescence in *E. coli* from ACE2NTD mutant libraries by simply screening for green colonies among thousands of transformants. It has been shown before that membrane protein fusion to GFP can create fusion problem for GFP with reduced fluorescence [43]. GFP-expressing cell suspensions and lysate (IPTG-induced) showed a yellow/green color under normal light (**Suppl. Fig. S2C, D**).

### Expression of MBP-RNase A (bovine) and MBP-RNase I fusions

A schematic diagram of the *E. coli* RNase I and bovine RNase A fusion with MBP is shown in **Fig. 2A**. The fusion proteins should be exported to the periplasm and the MBP signal peptide being cleaved, but some fusion proteins could exist in the cytoplasm due to the protein overexpression and overwhelming of the export system. Analysis of four cell lysates (total proteins) from NEB Express indicated a strong induced band of 55 to 60 kDa in three out of four samples, in close agreement with the predicted MBP-RNase A fusion (58.3 kDa) (**Fig. 2B**). The inserts in plasmid clones (#1, #3, and #4) were sequenced to confirm the correct DNA sequence. After centrifugation, most of the fusion protein was pelleted, and a small fraction was found in the supernatant (**Fig. 2C, D**). Attempt to transfer the MBP-RNase A expression clone into NEB SHuffle strain was not successful (few colonies were found in transformation, data not shown), suggesting the fusion protein may be toxic to the host. Bovine RNase A is a single-strand specific enzyme, but at high concentrations it can cleave dsRNA as well [44]. Its over-expression in T7 SHuffle cells appeared to be lethal to the host. A signal peptide may be added to bovine RNase A to be exported to periplasm for its over-expression in *E. coli*. Due to the low yield of soluble MBP-RNase A (bovine) fusion, the purification was not pursued further.

**Fig. 2.**
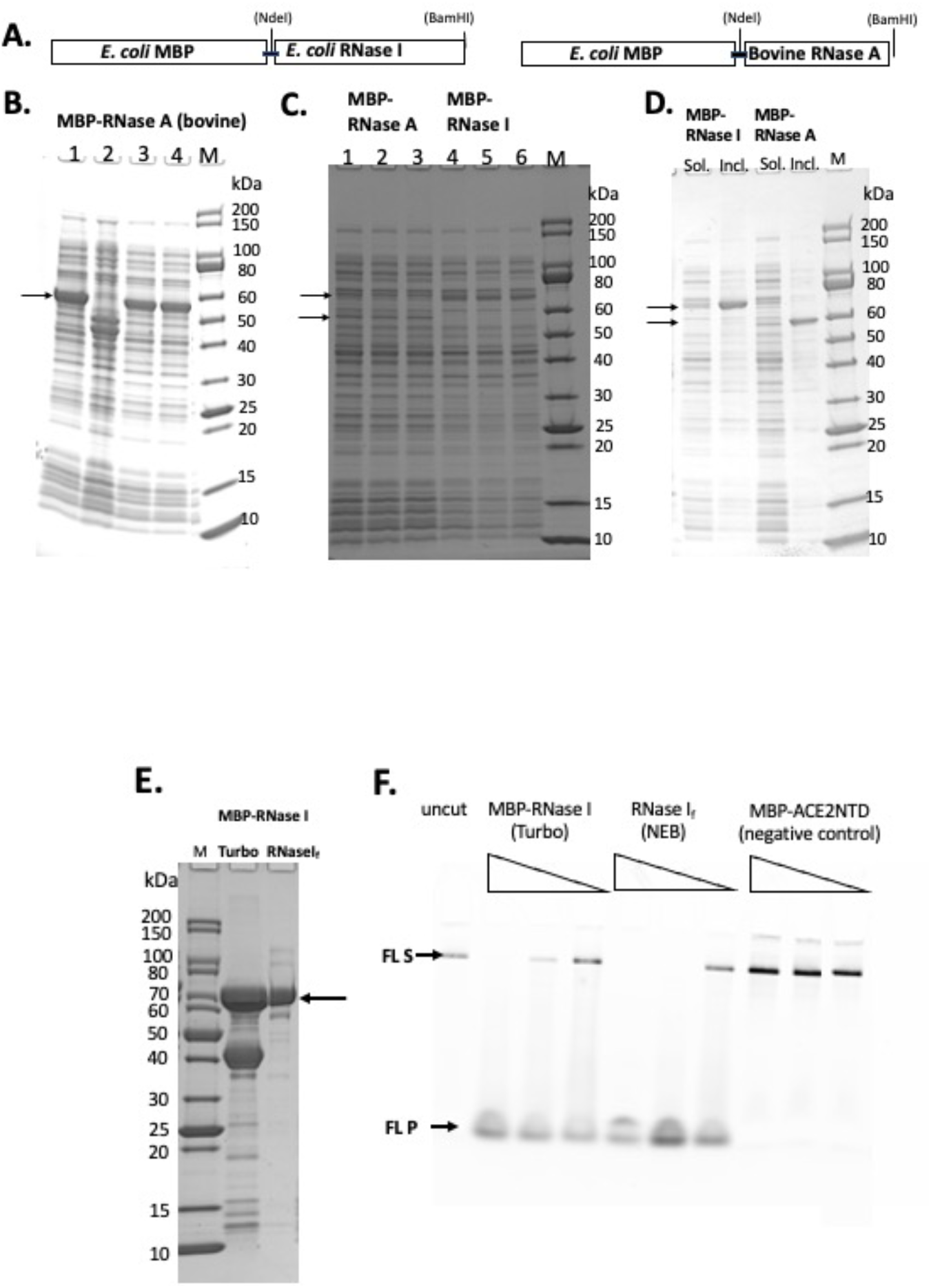
SDS-PAGE analysis of MBP-RNase I and MBP-RNase A fusions. **A.** A schematic diagram of MBP-RNase A fusion. **B.** Total cellular proteins from IPTG-induced cells (NEB Express). **C.** Supernatant (soluble) of the fusion proteins (3 independent isolates each). Arrows indicate the fusions. **D.** Comparison of supernatant (Sol.) and pellet (Inc.) of the fusions. M, protein size marker (NEB). **E.** SDS-PAGE analysis of partially purified MBP-RNase I fusion. Lane 1, protein size marker; lane 2, MBP-RNase I fusion purified from IPTG-induced NEB Turbo cells; lane 3, RNase I_f_ (NEB). **F.** RNase activity on Fluorescein-labeled RNA (300 nt). No enzyme (uncut) and MBP-ACE2NTD fusion serve as negative controls.

A PCR fragment encoding the *E. coli* RNase I was inserted into the pMAL-p5x vector to construct MBP-RNase I fusion. Analysis of 11 cell lysates (total proteins) from IPTG-induced cells (NEB Express) indicated a strong induced band of ~70 kDa in 8 out of 11 samples, in agreement with the predicted MW (71.7 kDa) (**Fig. 2C, Suppl. Fig. S3**). After centrifugation, most of the fusion protein was found in the pellet, and a small fraction remained in the supernatant (**Fig. 2C, D**). Attempt to express the fusion in T7 SHuffle cells did not improve the expression level (data not shown). MBP-RNase I fusion appeared to be toxic to T7 SHuffle host since only low-density cells could be obtained in liquid culture at 37°C. The protein yield of MBP-RNase I was highest in NEB Turbo cells, probably due to the robust cell growth and higher cell weight per liter culture. **Fig. 2E** shows the partially purified MBP-RNase I fusion in comparison with RNase I_f_, (RNase I_f_ = MBP-RNase I fusion from NEB as a research grade ribonuclease, M0243S). The enzyme preparation also contained the host MBP. In order to eliminate the MBP, it will be necessary to use an MBP-deficient strain as the expression host. In a ribonuclease activity assay, the partially purified MBP-RNase I is active in digestion of a RNase substrate (fluorescein-labeled 300 nt RNA) (**Fig. 2F**).

Previous studies showed that *E. coli* RNase I does not require divalent cations for nuclease activity, and the other eight ribonucleases from *E. coli* are only active in the presence of divalent cations [37]. To demonstrate the ribonuclease activity for the partially purified MBP-RNase I, we incubated a low range ssRNA ladder (N0364S, NEB) with the enzyme (3 enzyme dilutions at 2 μg, 0.2 μg, and 0.02 μg) in a high salt buffer supplemented with EDTA, Ni^2+^, Ca^2+^, Mn^2+^, Co^2+^, or Mg^2+^. MBP-RNase I is most active in Ca^2+^, Mn^2+^, and Co^2+^ buffers. It is also active in EDTA, Ni^2+^, and Mg^2+^ buffers, but with limited digestion at low enzyme input (**Suppl. Fig. S4A**). It is possible that in Ca^2+^, Mn^2+^, and Co^2+^ buffers, MBP-RNase I activity is stimulated at the low enzyme concentration. It was concluded that MBP-RNase I fusion can be expressed at moderate level in NEB Turbo and NEB Express cells. A small fraction of the fusion protein was found in the soluble fraction and can be purified by chromatography through amylose column. The partially purified fusion is active in digestion of RNA in the absence of divalent cations or in the presence of EDTA.

### Expression and purification of E. coli RNase I (6xHis)

To improve the expression level of RNase I, we also expressed a C-terminal 6xHis-tagged version. *E. coli rna* gene was inserted into the T7 expression vector pET21b. Its expression level was compared in two T7 expression strains (T7 Express and Nico (λDE3). *E. coli* RNase I has its own signal peptide to be exported to periplasm. But we detected two versions in over-expression: RNase I precursor with the signal peptide in cytoplasm and the mature enzyme without the signal peptide. The proteins eluted from C2529 cell lysate contained fewer contaminating proteins than C2566 cells, due to the knock-out mutations of a few Histidine-rich proteins in C2529 (**Fig. 3A**). Two versions of RNase I appeared to be co-purified: a longer cytoplasmic version (cRNase I) with the signal peptide (precursor, 30 kDa), and the shorter version of the periplasmic form (27 kDa). The partially purified enzyme is active in digestion of FAM-labeled COVID-19 RNA (60 mer) in a high salt buffer in the absence of MgCl_2_. We also tested the ribonuclease activity in the presence and absence of divalent cations. RNase I (6xHis) is active in the presence of 10 mM EDTA. The RNase activity is mostly active in buffers with Ca^2+^, Mn^2+^, or Co^2+^ at low enzyme concentration (data not shown).

**Fig. 3.**
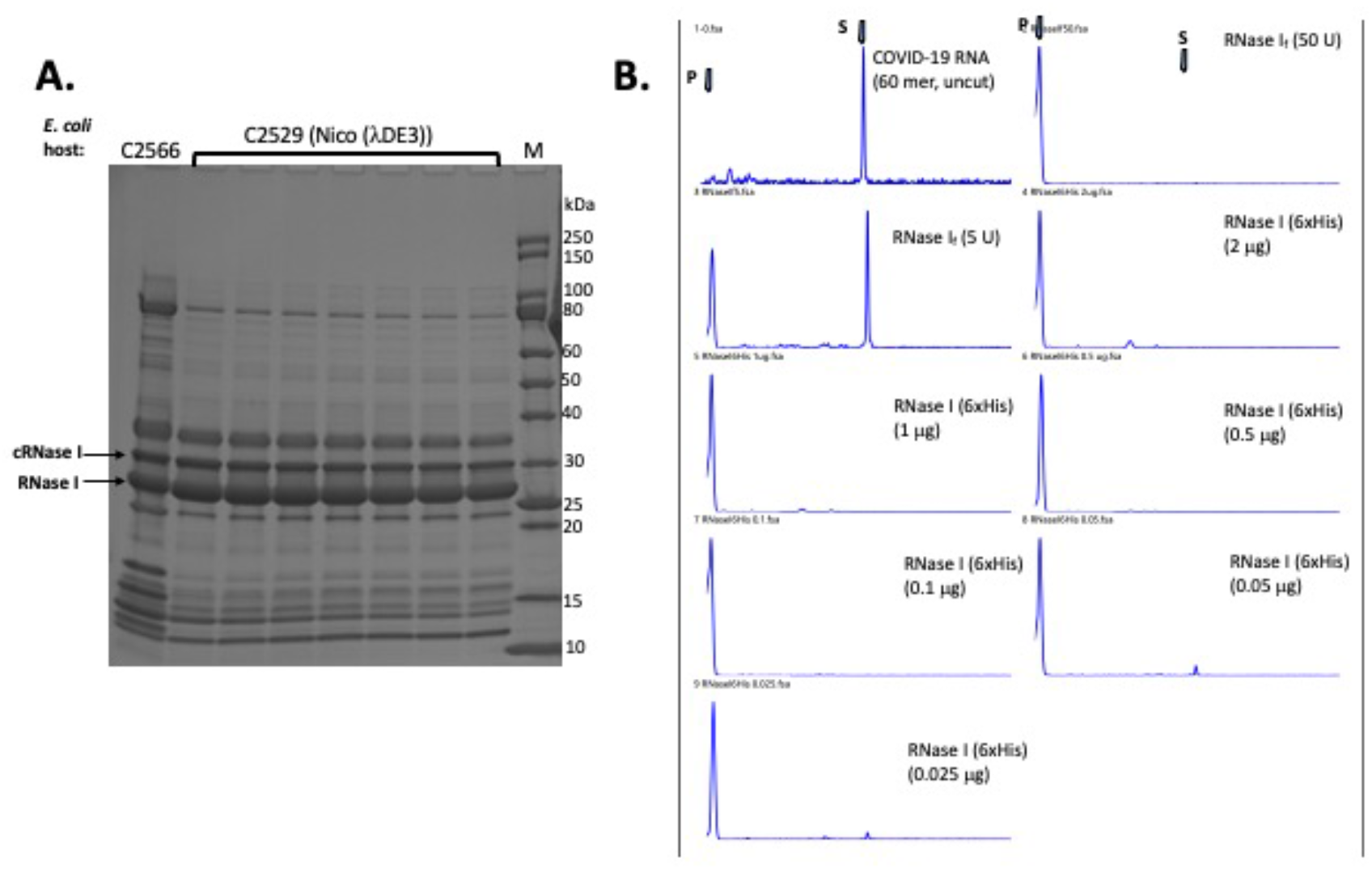
Purification of RNase I (6xHis) and RNase activity assay. **A.** Purification of RNase I (6xHis) from Nickel-NTA agarose column. Lane 1, RNase I (6xHis) pooled fractions from a nickel column (purified from T7 Express cell extract). Arrows indicate the cytoplasmic RNase I precursor (cRNase I) with the signal peptide (predicted MW 30.7 kDa), and the periplasmic RNase I with the signal peptide removed (predicted MW 27.0 kDa). **B.** RNase activity on a FAM-labeled COVID-19 RNA (60 mer). S = substrate; P = cleavage product(s). Positive controls, 50 and 5 U of RNase I_f_ (MBP-RNase I fusion, NEB). RNase I (6xHis) enzyme titration (2 μg to 25 ng protein) was used in the activity assay to digest fixed amount of RNA (16 nM) in NEB buffer 3 at 37°C for 1 h. Proteinase K (1.6 U) was added to remove RNase I. The final cleavage products were analyzed by capillary electrophoresis (CE) and peaks were visualized by PeakScan.

### Expression and purification of E. coli RNase III (6xHis)

Since the SARS-CoV-2 RNA genome also contains regions of dsRNA (e.g. RNA hairpin stem loops structures, frame shift element) (reference: RJ Andrews et al. (bioRxiv. Preprint. 2020 Apr 18. doi: 10.1101/2020.04.17.045161), RNase III may be needed to digest the duplex RNA sequences. The *E. coli rnc* gene encoding RNase III was cloned into pET21b to generate a C-terminal 6xHis-tagged protein. RNase III expression level after IPTG induction was examined in three T7 expression strains: T7 SHuffle (C3026), T7 Express with LysY and *lacI^q^* (C3013) and Nico (λDE3) (C2529). **Fig. 4A** shows the expression levels (soluble fractions/supernatant) in the three strains. RNase III (6xHis) was partially purified from a Nickel-NTA agarose column or Ni magnetic beads (**Fig. 4B**). It is active in digestion of a 40mer RNA duplex and dsRNA ladder. The 40mer duplex RNA and dsRNA ladder were digested into smaller fragments as detected in the agarose gel (**Fig. 4C)**. In a control experiment, the ShortCut RNase III (MBP-RNase III fusion) also digested the RNA substrates and gave rise to similar shortened fragments. It was concluded that RNase III (6xHis) can be expressed in *E. coli* at high level and it is not toxic to *E. coli* T7 expression strains. The expression level is comparable in the three strains tested.

**Fig. 4.**
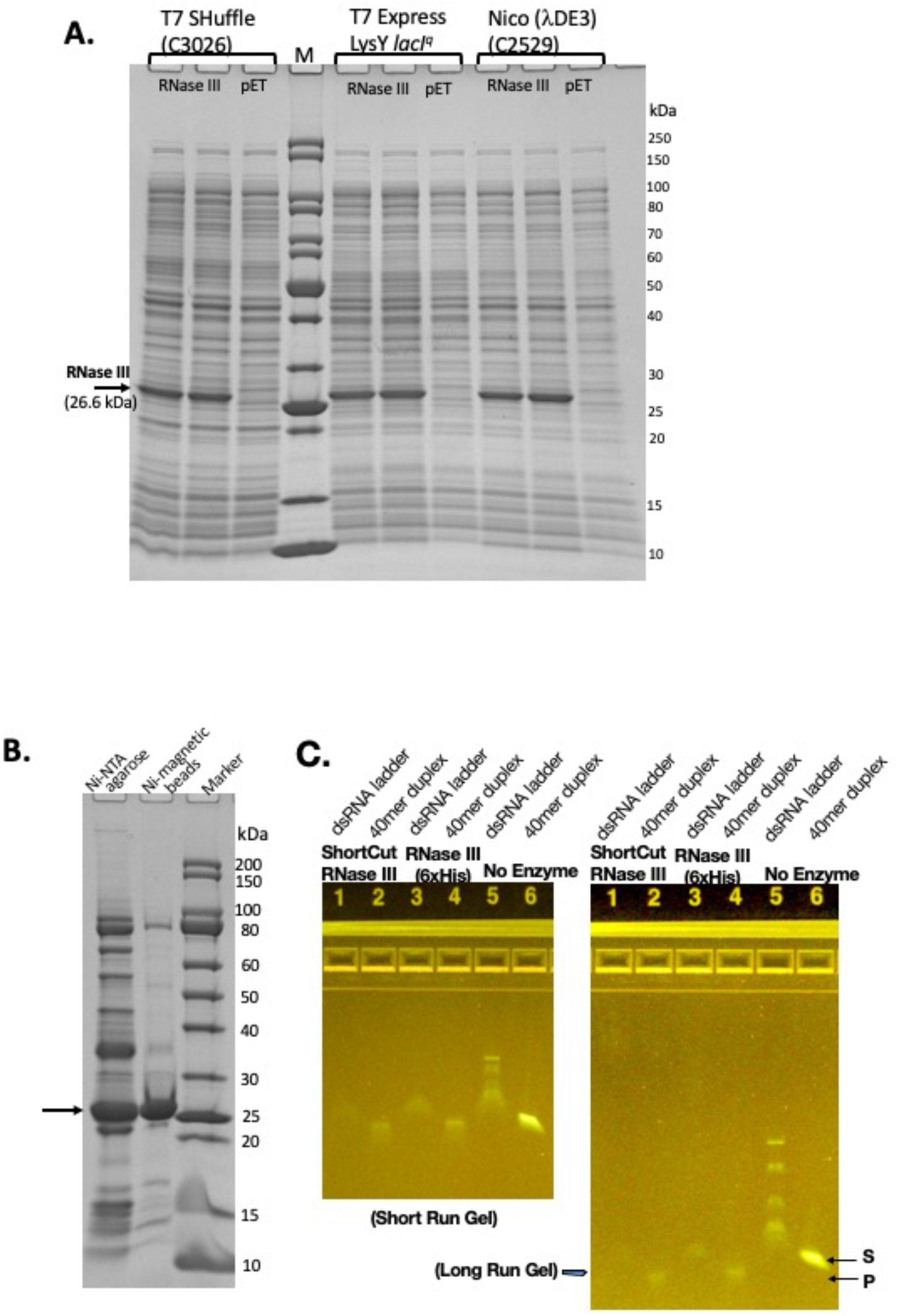
Expression and purification of *E. coli* RNase III and RNase activity assay on dsRNA. **A.** RNase III expression level in three *E. coli* T7 strains: T7 Shuffle (C3026), T7 Express with *lacI^q^* and LysY (C3013), and Nico (λDE3). **B.** Purified RNase III from nickel-NTA agarose column chromatography and Ni magnetic beads. **C.** Ribonuclease activity assay on dsRNA.

### Expression and purification of RNase I-ACE2 (6xHis) fusion protein

Since our constructs of MBP-ACE2NTD and RNases were shown to be folded and soluble, we next sought to generate the fusion protein that can bind S protein and exhibit cleavage activity. The RNase I-ACE2NTD (6xHis) fusion was expressed in Nico (λDE3). Most of the fusion was found in the inclusion body and only a small fraction was found in the supernatant as detected by anti-6xHis Ab or anti-ACE2 monoclonal Ab in Western blot and SDS-PAGE (**Fig. 5A and B**, **Suppl. Fig. S5**). The RNase I-ACE2NTD (6xHis) fusion pellet was subjected to a refolding procedure and further purified by binding to Ni magnetic beads or Ni spin column (**Fig. 5C**). The refolded fusion protein is active in digestion of a 300-nt RNA substrate (**Fig. 5D**), similar to the positive controls RNase I (6xHis) and MBP-RNase I. The fusion enzyme purified through Ni magnetic beads and spin column is also active in digestion of COVID-19 RNA (60 mer) (**Fig. 5E**). We have not tested the ribonuclease activity on full-length COVID-19 RNA due to local bio-safety regulation on the virus.

**Fig. 5.**
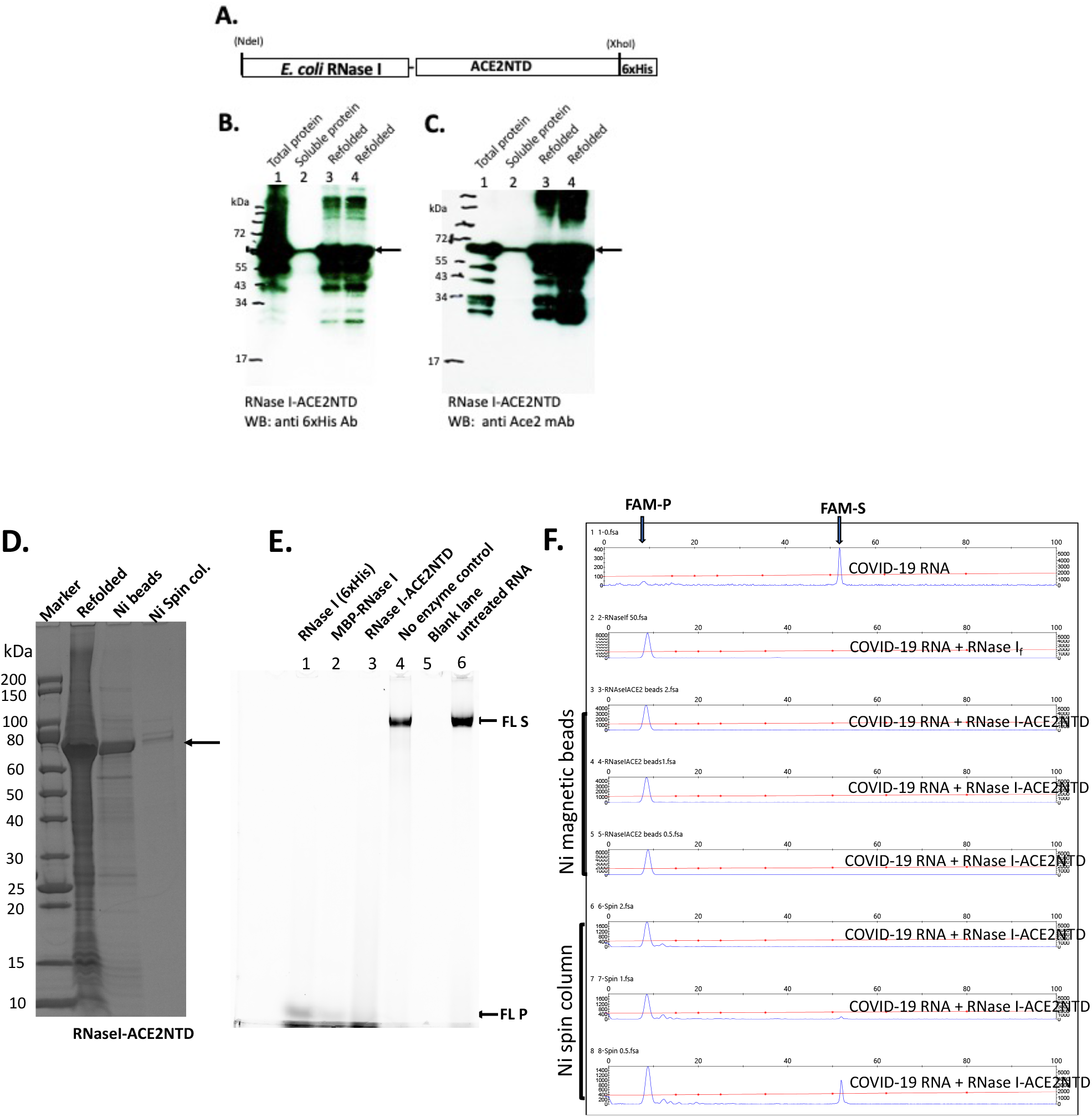
SDS-PAGE and Western blot analysis of RNase I-ACE2NTD fusion and activity assays. **A.** Schematic diagram of RNase I-ACENTD (6xHis) fusion. **B.** Western blot analysis of RNase I-ACE2NTD in total protein, supernatant (soluble), and refolded protein using anti-6xHis Ab. **C.** Same as in **B**, except using anti-ACE2 monoclonal Ab. **D.** SDS-PAGE analysis of the refolded RNase I-ACE2NTD fusion and further purified protein by Ni magnetic beads and Ni spin column. **E.** RNase I-ACE2NTD (refolded) ribonuclease activity on Fluorescein (FL)-labeled DNA (300 nt) in NEB buffer 3. RNase I (6xHis) and MBP-RNase I were used as positive controls. **F.** Ribonuclease activity of RNase I-ACE2NTD (purified by Ni magnetic beads or Ni spin column) on COVID-19 RNA (60mer). RNase I_f_, a positive control. FAM-S, FAM-labeled substrate; FAM-P, FAM labeled cleavage product(s).

### Expression and purification of hRNase A-ACE2NTD150 fusion with and without export signal peptide

A schematic diagram of fusion of hRNase A with ACE2NTD150 (a shorter version of ACE2NTD) is shown in **Fig. 6A**. A short linker (GSAGSA) is inserted between the fusion partners. hRNase A’s native signal peptide was deleted and MBP signal peptide was added. The second clone does not carry any signal peptide (cytoplasmic version). The over-expressed fusions could be detected in IPTG-induced cell lysate (total proteins) as shown in **Fig. 6B**. The two fusions appeared as closely migrated doublet. It may have something to do with multiple disulfide bonds in hRNase A (8 Cys residues in total, potentially forming 4 disulfide bonds). With protein loading dye, DTT and heating before SDS-PAGE, some of the disulfide bonds may be reduced to -SH groups, creating a mixture of isoforms with different secondary structures (S-S vs -SH). Another possibility is that export systems are overwhelmed and some of the protein with signal peptide is not secreted. To confirm the identity of the engineered fusion, we perform Western blot using mouse anti-His Ab and anti-ACE2 monoclonal Ab. **Fig. 6C** shows the soluble fusion proteins detected by the antibodies. There are some minor degradation products from the fusions as detected by the anti-ACE2 Ab. We also tested fusion expression level in Nico (λDE3) and NEB SHuffle T7 (B strain). hRNase A-ACE2NTD150 fusions with or without the MBP signal peptide can be over-expressed in Nico (λDE3) cells (data not shown). However, hRNase A-ACE2NTD150 fusion expression is toxic to the Shuffle strain: hRNase A-ACE2NTD150 with signal peptide formed small colonies in transformation of T7 SHuffle. No transformants were found with plasmid carrying the coding sequence for hRNase A-ACE2NTD150 (ΔS, lacking MBP signal peptide). This result is similar to the toxic effect by MBP-RNase A (bovine) in transformation of T7 SHuffle (K strain).

**Fig. 6.**
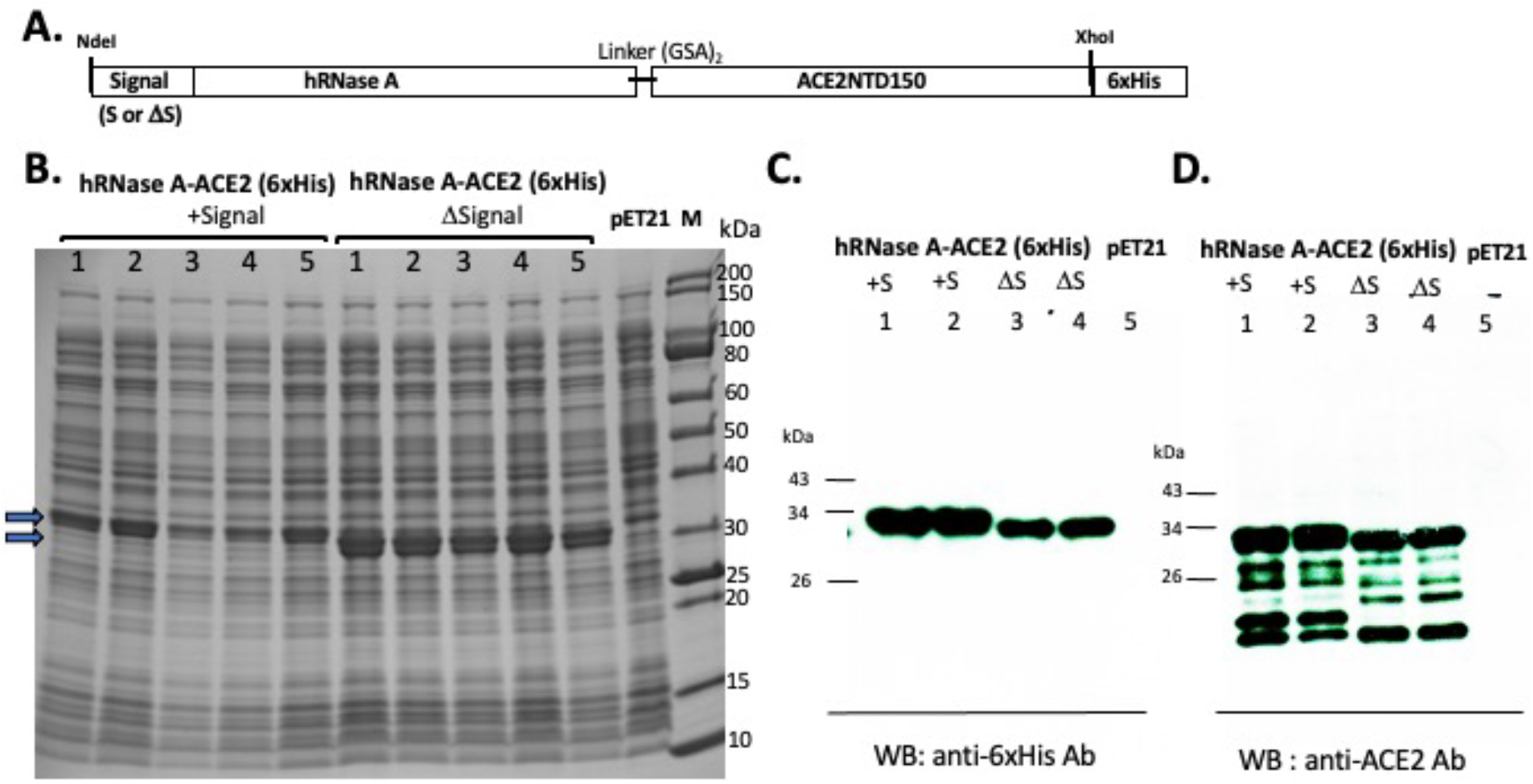
Expression of hRNase A-ACE2NTD150 in T7 Express *LysY/lacI^q^* (C3013). **A.** A schematic diagram of the fusion protein. **B.** SDS-PAGE analysis of five isolates of hRNase A-ACE2NTD150 (five isolates with MBP signal peptide, five clones without MBP signal peptide). Total cell lysate of pET21b serves as a negative control. **C and D.** Western blot analysis of the fusion proteins (soluble fraction/supernatant) by anti-6xHis Ab and anti-ACE2 monoclonal Ab.

Most of the hRNase A-ACE2NTD150 fusions (with or without MBP signal peptide) were found in the pellet following centrifugation (data not shown). However, there are still some soluble fusions in the supernatant. The fusion protein hRNase A-ACE2NTD150 (6xHis) (no signal peptide) was partially purified by chromatography through a Ni-NTA agarose column or binding to Ni magnetic beads (**Fig. 7A**). The purified fusion proteins were active in cleaving FAM-labeled COVID-19 RNA (60mer) (**Fig. 7B**). The substrate RNA completely disappeared after 30 min digestion (at 0.1 μg to 2 μg of the fusion protein). Both RNA substrates and cleavage products were detected after digestion at the lower enzyme concentration (at 1.25 to 10 ng) (**Fig. 7B**). Some smaller cleavage products less than 10 nt were not easily detected in the CE assay. To further confirm the ribonuclease activity, we digested a 300-nt RNA and a ssRNA ladder (low MW). Both RNA substrates were degraded by the fusion enzyme as detected on a 6% PAG-urea gel and staining with SYBR green (**Fig. 7C**). It was concluded that the fusion enzyme hRNase A-ACE2NTD150 displays ribonuclease activity on both 60mer COVID-19 RNA and other RNA substrates ranging from 50-1000 nt. We have not tested ribonuclease activity on the full-length SARS-CoV-2 viral RNA. The binding affinity to RBD domain of the Spike protein remains to be tested.

**Fig. 7.**
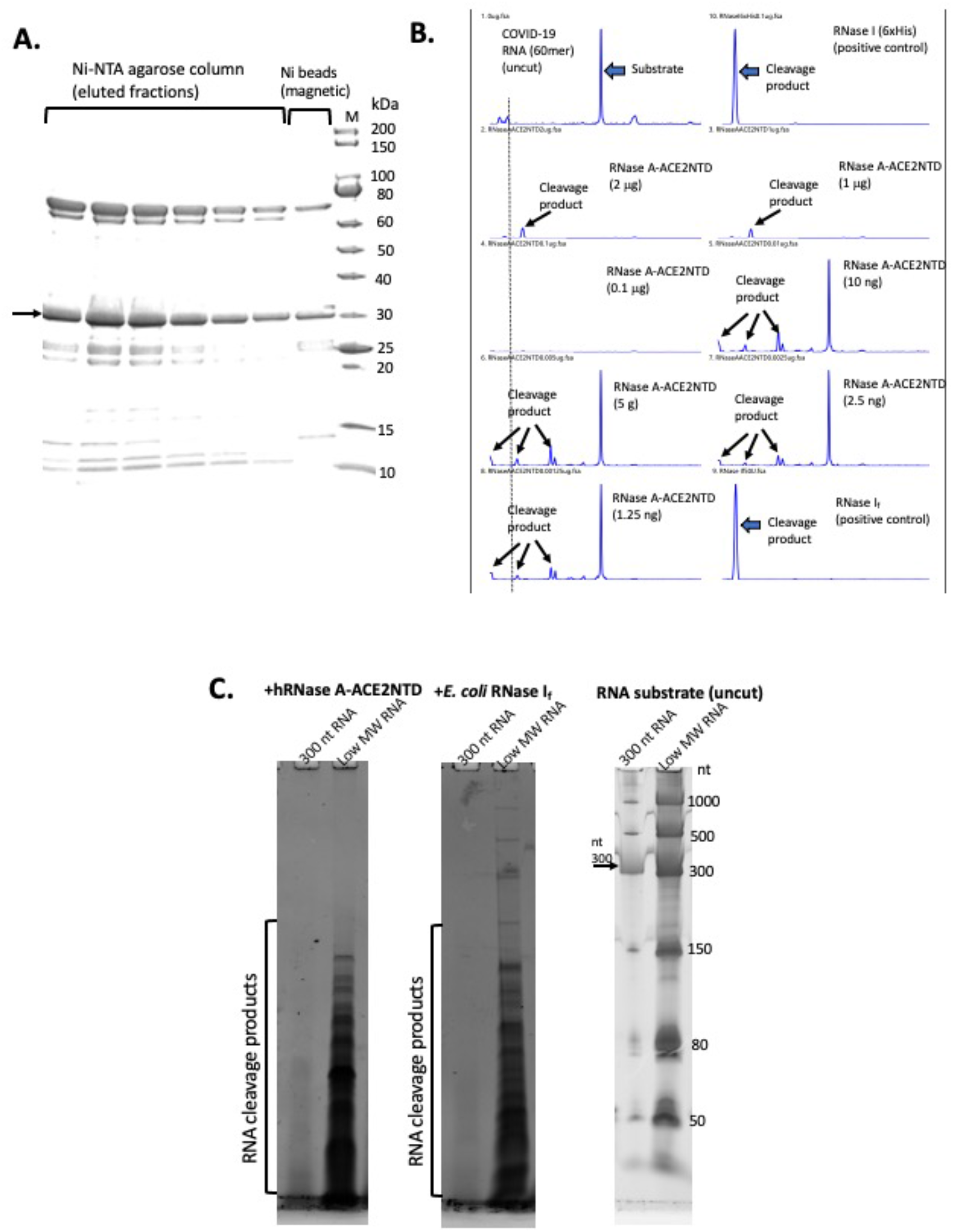
SDS-PAGE analysis of purified hRNase A-ACE2NTD150 (6xHis) and ribonuclease activity assays. **A.** Partially purified hRNase A-ACE2NTD150 (6xHis) (no signal peptide) by Ni-NTA agarose column chromatography or by binding to Ni magnetic beads. **B.** RNase activity on FAM-labeled COVID-19 RNA (60mer). Arrows indicate the substrate and cleavage products in the CE assay. **C.** RNase activity assay on a 300-nt RNA and low MW RNA ladder as analyzed on a 6% PAG-urea gel (stained with SYBR green and visualized on a Typhoon Imager).

The fusion protein expression and purification conditions and activity assays on RNA substrates are summarized in **Table 1.**

**Table 1.** *E. coli* expression strains, purification method, enzyme activity/potential usage.

| **Protein construct** | ***E. coli* host** | **Purification method** | **Protein function or RNase activity** |
| --- | --- | --- | --- |
| MBP-ACE2NTD<sup>a</sup> | T7 SHuffle or NEB Express | Amylose column or protein refolding, amylose magnetic beads | Binding to S-RBD |
| MBP-RNase I<sup>b</sup> | NEB Turbo or NEB Express | Amylose column | Cleaving 300-nt RNA |
| ACE2NTD-GFP | T7 SHuffle | Not determined (ND) | Low green fluorescence |
| RNase I (6xHis) | T7 Express (LysY/*lacI*<sup>q</sup>) or Nico (λDE3) | Ni-NTA agarose column | Cleaving RNA ladder and 60 mer COVID-19 RNA |
| RNase III (6xHis) | T7 Express (LysY/*lacI*<sup>q</sup>) or Nico (λDE3) | Ni-NTA agarose column | Cleaving duplex RNA and dsRNA ladder |
| RNase I-ACE2NTD (6xHis) | Nico (λDE3) or BL21 (λDE3) | Protein refolding and Ni magnetic beads | Cleaving 300-nt RNA and 60 mer COVID-19 RNA |
| hRNase A-ACE2NTD150 (6xHis)<sup>c</sup> | Nico (λDE3) or T7 Express (LysY/*lacI*<sup>q</sup>) | Ni-NTA agarose, Ni magnetic beads | Cleaving COVID-19 RNA and ssRNA ladder |

- a. ACE2 signal peptide removed. - b. Native RNase I signal peptide removed. - c. Human RNase A original signal peptide removed.

## Discussion

This work demonstrated that MBP-ACE2NTD, MBP-RNase I, RNase I (6xHis), RNase III (6xHis), RNase I-ACE2NTD (6xHis), hRNase A-ACE2NTD150 can be expressed in *E. coli* either as soluble protein or refolded protein from inclusion bodies. We tested a number of *E. coli* expression strains and found that MBP-ACE2NTD expression was partially soluble in T7 SHuffle strain, while RNase I-ACE2NTD (6xHis) fusion was best expressed in Nico (λDE3) or BL21 (λDE3) as refolded protein from inclusion body. RNase I (6xHis) and RNase III (6xHis) expressions were more comparable among the T7 expression strains tested in this work. The over-expression of MBP-RNase A (bovine) hRNase A-ACE2NTD150 fusions, however, is toxic to T7 SHuffle strains. We conclude from this work that different *E. coli* expression strains need to be tested in order to optimize the expression condition. We only tested low temperature induction at 16°C to 18°C and the protein induction level at higher temperatures has not been examined. It is anticipated that protein expression level may be increased due to the higher cell weight at the higher temperature with possible trade-off of lower protein solubility. Large-scale fermentation and low-cost purification will be required for mass production and use of these enzymes as anti-viral agents. Other protein expression system, for example, the yeast expression system could be used to over-express human ACE2 receptor and TMPRSS2. In addition, human ACE2 receptor protein and the serine protease can be expressed and exported to the surface of bacteria and yeast for cell surface display [45]. Homing endonuclease variants have been expressed and selected successfully in yeast surface display; the engineered endonuclease mutants displayed altered specificities with a range of cleavage activities *in vitro* [46]. The fusion of ACE2NTD to *E. coli* FimH protein to display on the cell surface did not generate positive results in our preliminary study: the fusion coding sequence can be constructed in DNA level, but no fusion protein was detected by anti-6xHis Ab to detect FimH signal peptide-ACE2NTD-FimH (6xHis) fusion. Presumably the fusion is toxic to the host cell.

The ACE2NTD-GFP fusion may be used as a bio-sensor if more soluble ACE2NTD variants could be isolated. The tight binding of COVID-19 variants to the ACE2NTD-GFP fusion may alter the fluorescence signal intensity.

Future research may be directed towards making full-length recombinant ACE2 receptor from *E. coli* with co-expression of chaperon proteins to increase solubility (chaperone plasmid set, Takara Bio)[47]. In a previous work, we found co-expression of *E. coli* GroEL/ES proteins with GmrSD endonuclease greatly improved the solubility of GmrSD [48]. To reduce protein purification cost, expression of RNase I-ACE2 (6xHis) and hRNase A-ACE2 (6xHis) fusions may be carried out using the *K. lactis* yeast expression system (NEB) to export the recombinant proteins into the culture medium for industrial production of the fusions.

Other SARS-CoV-2 structure proteins and human membrane proteins (e.g. TMPRSS2 serine protease catalytic domain) or furin protease may be added to the recombinant protein mixture to trigger virus binding and viral RNA release (TMPRSS2 and Furin cleave SARS-CoV-2 S protein into S1 and S2 domains). Use of mild detergent, EDTA, and proteases may also destabilize the infectious virus particles and partial release of viral RNA for destruction. More research is urgently needed to validate the enzymatic approach to reduce virus counts and transmission in air droplets and to combat highly transmissible COVID-19 variants.

It is noted that the experiments described in this work were carried out with small fragments of RNA *in vitro* and that work with real virus is needed to fully evaluate the effectiveness as an anti-viral strategy. We are actively seeking collaborations with academic labs and government institutes.

## Acknowledgement

This work was supported by New England Biolabs, Inc. We thank Andy Gardner, Rich Roberts and Tom Evans for their support and idea development, Andy Gardner for critical comments, Sebastian Gruenberg and Dongxian Yue for research materials, Nono Go for the artistic illustration, and DNA sequencing core lab (NEB) for sequencing and CE analysis.

## Supplementary Material: Suppl. Fig. S1-S5

**Suppl. Fig. S1.**
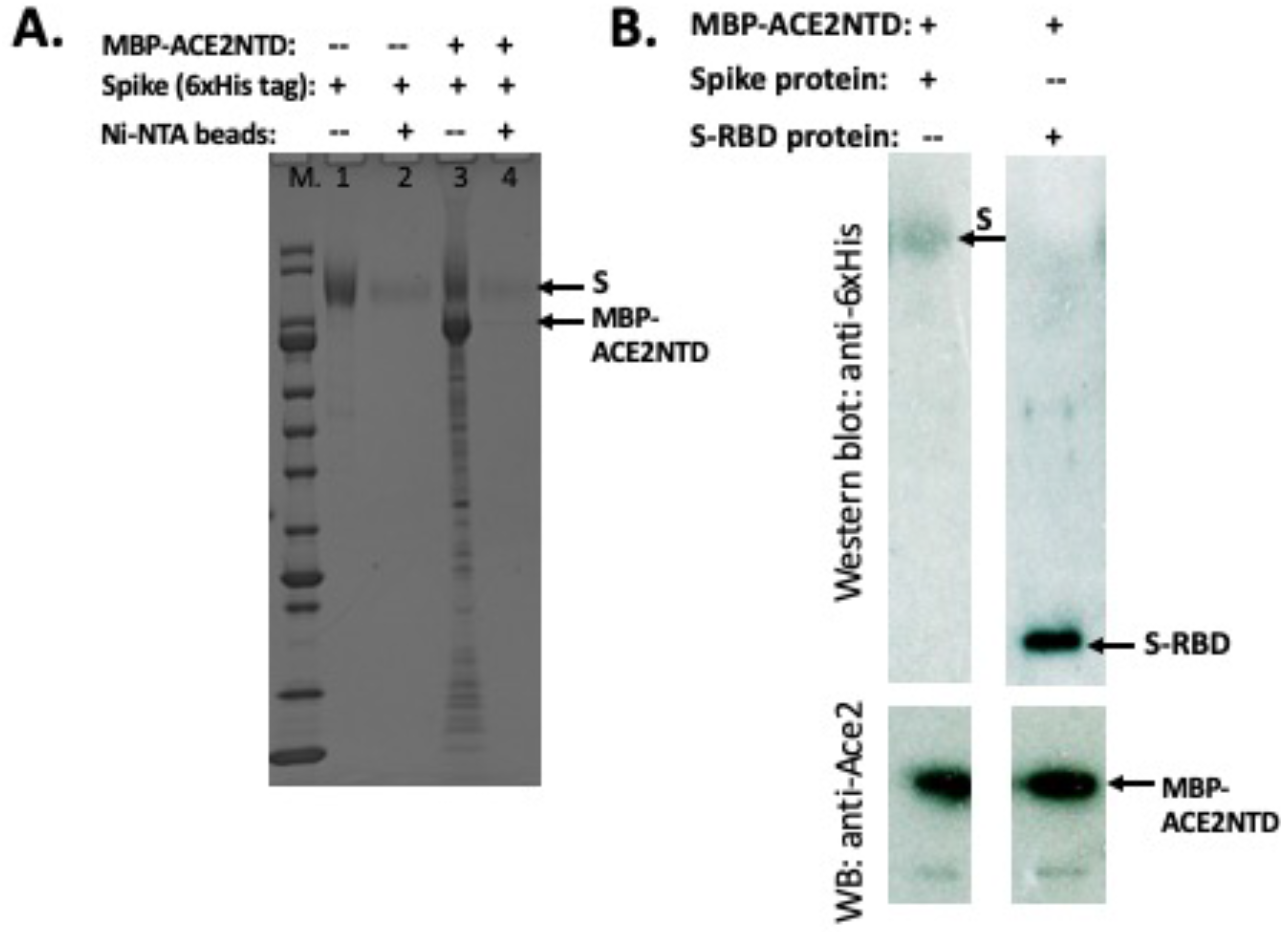
Protein pull-down assays for MBP-ACE2NTD, Spike (S) and RBD protein. **A.** His-tagged Spike protein bound to Ni magnetic beads incubated with MBP-ACE2NTD and co-eluted proteins. **B.** MBP-ACE2NTD bound to amylose magnetic beads incubated with Spike or RBD protein and co-eluted proteins as detected by anti-6xHis and anti-ACE2 Abs.

**Suppl. Fig. S2.**
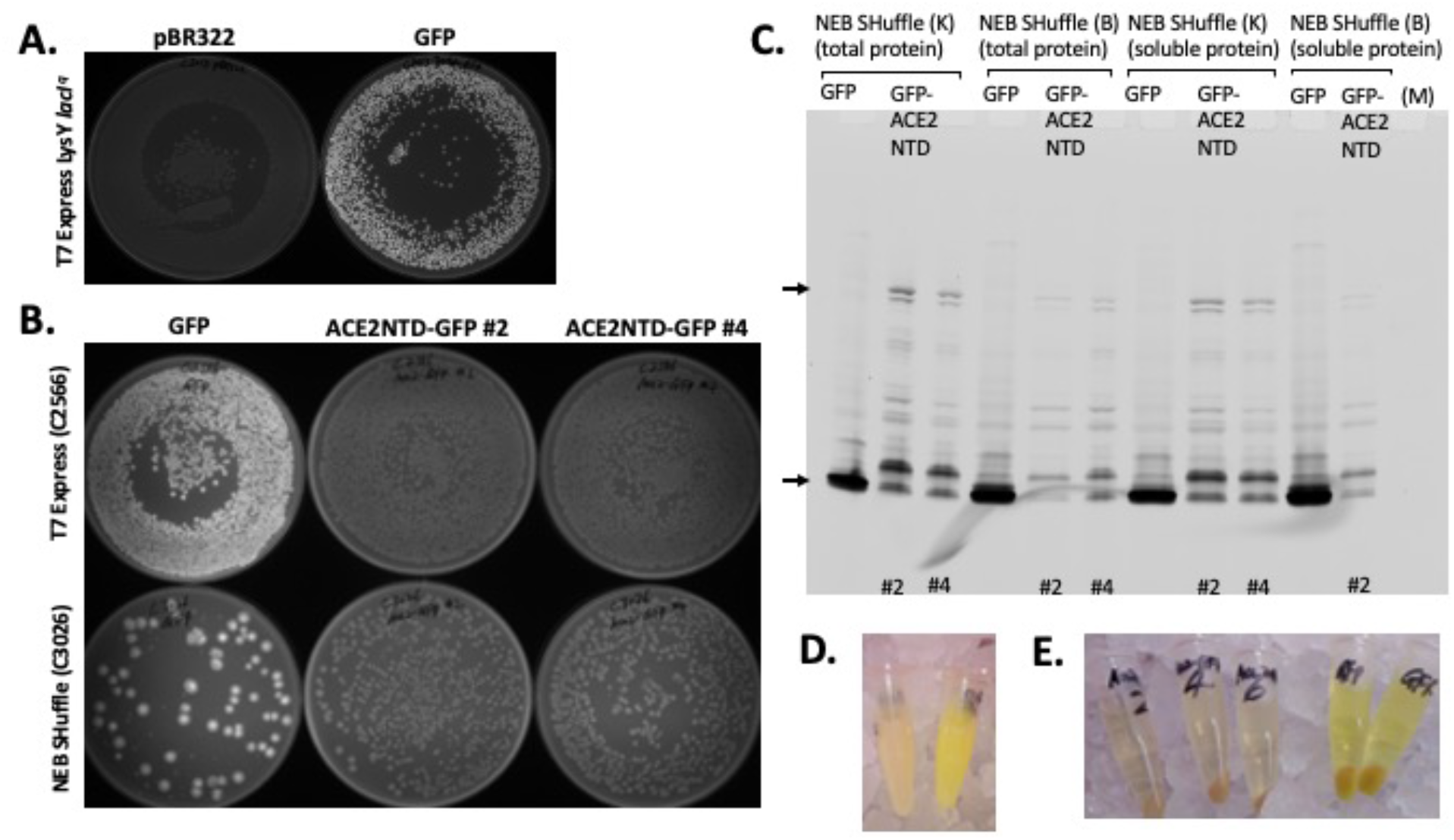
Green fluorescence detection for GFP and ACE2NTD-GFP fusion protein. **A.** T7 Express *(LysY/lacI^q^) cells carrying pBR322 (a negative control) and pBR322-P_T5_-*lacO*-DasherGFP visualized under long UV light.* **B.** *E. coli* colonies (Amp^R^ transformants) of NEB SHuffle and T7 Express cells expressing GFP or ACE2NTD-GFP fusion visualized under long UV light. GFP expressing colonies show bright green color. T7 Express [ACE2NTD-GFP] shows no green fluorescence. NEB SHuffle [ACE2NTD-GFP] shows a weak green fluorescence. **C.** Total proteins from cell lysates of NEB SHuffle [ACE2NTD-GFP] (K and B strains) detected by fluorescence imaging at 520 nm (Cy2 channel). Arrows indicate GFP and ACE2NTD-GFP fusion, respectively. The Protein ladder shows no fluorescence and was not detected in the imager. **D.** Cells suspension of C3013 [pBR322] and C3013 [GFP]. **E.** T7 Express (C2566) cell lysates of ACE2NTD-GFP fusion and GFP. GFP expressing cells and cell lysate show yellow/green color under normal light as visualized by naked eyes.

**Suppl. Fig. S3.**
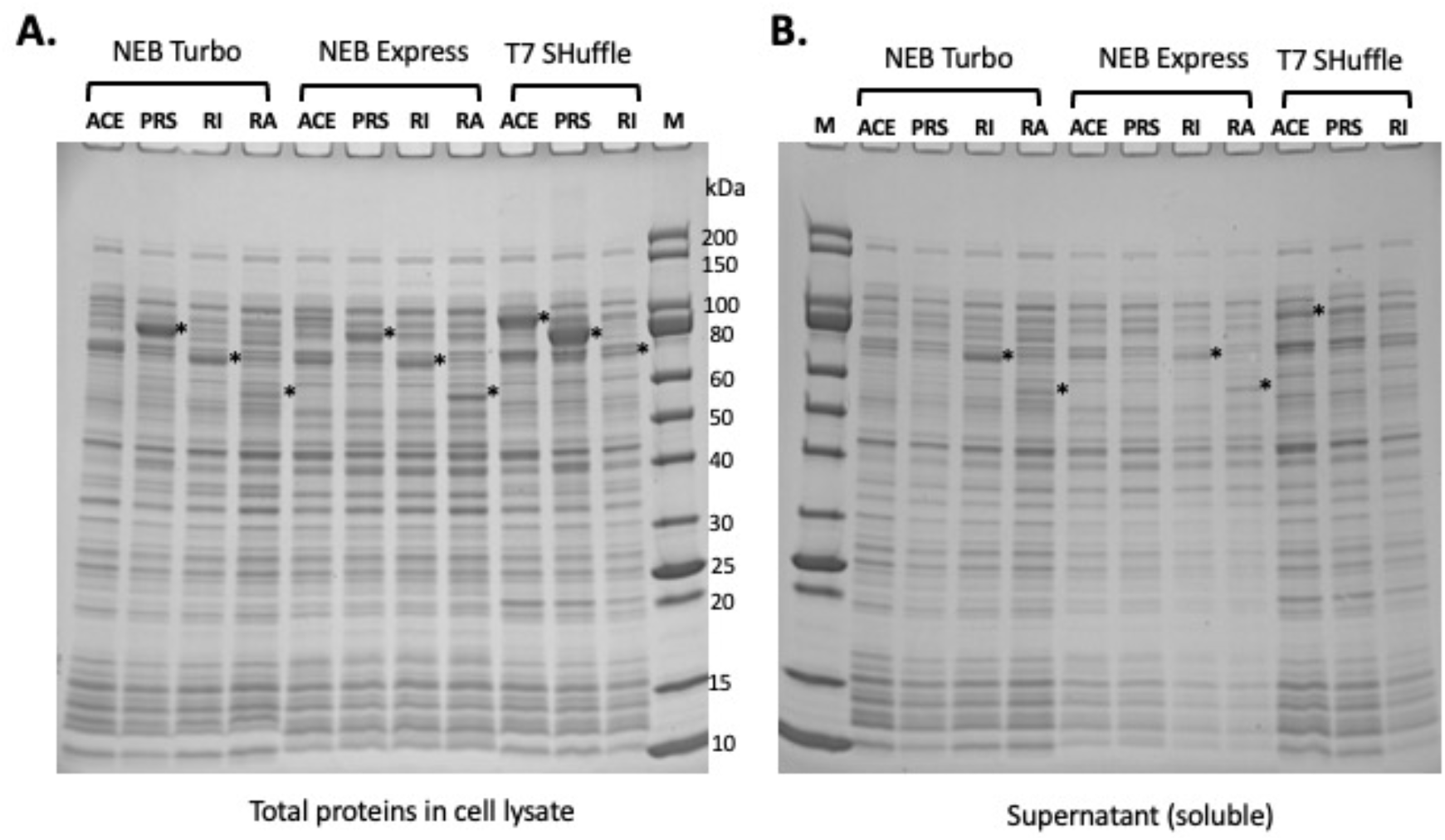
Comparison of protein expression in three *E. coli* strains: NEB Turbo, NEB Express, T7 SHuffle (K strain). MBP-ACE2NTD (ACE), MBP-TMPRSS2 (PRS, lacking the transmembrane domain), MBP-RNase I (RI), MBP-RNase A (RA). **A.** SDS-PAGE analysis of total proteins in cell lysate. **B.** SDS-PAGE analysis of soluble proteins (supernatant) in cell lysate. “*” indicates the expected target protein.

**Suppl. Fig. S4.**
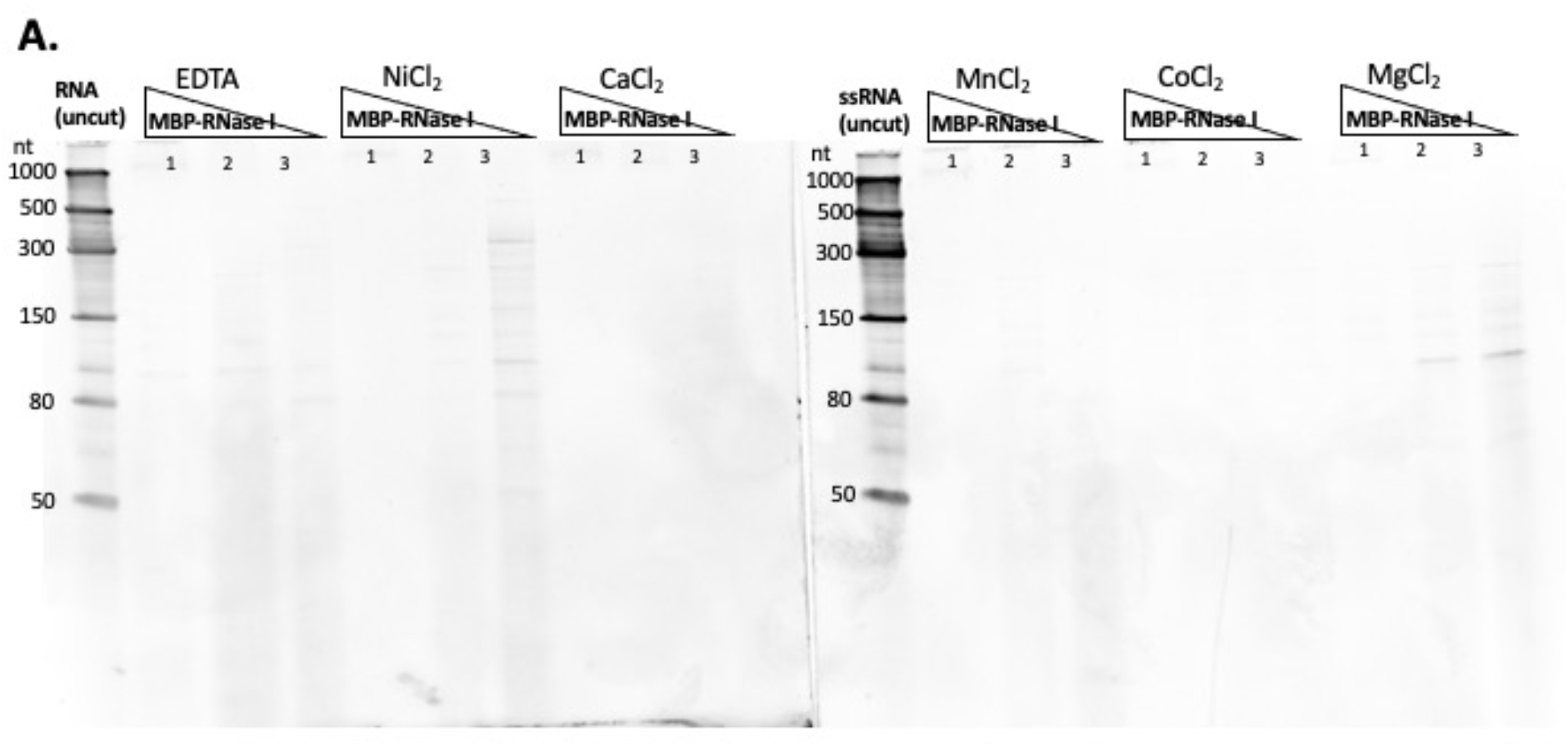
Ribonuclease activity in the presence of EDTA and divalent cations. **A.** MBP-RNase I ribonuclease activity assay. The low range ssRNA ladder (50 to 1000 nt long, NEB) was used as the substrate for RNase activity assay in a high sale buffer (100 mM NaCl, 50 mM Tris-HCl, pH 7.5) supplemented with divalent cations (1 mM) or EDTA (10 mM).

**Suppl. Fig. S5.**
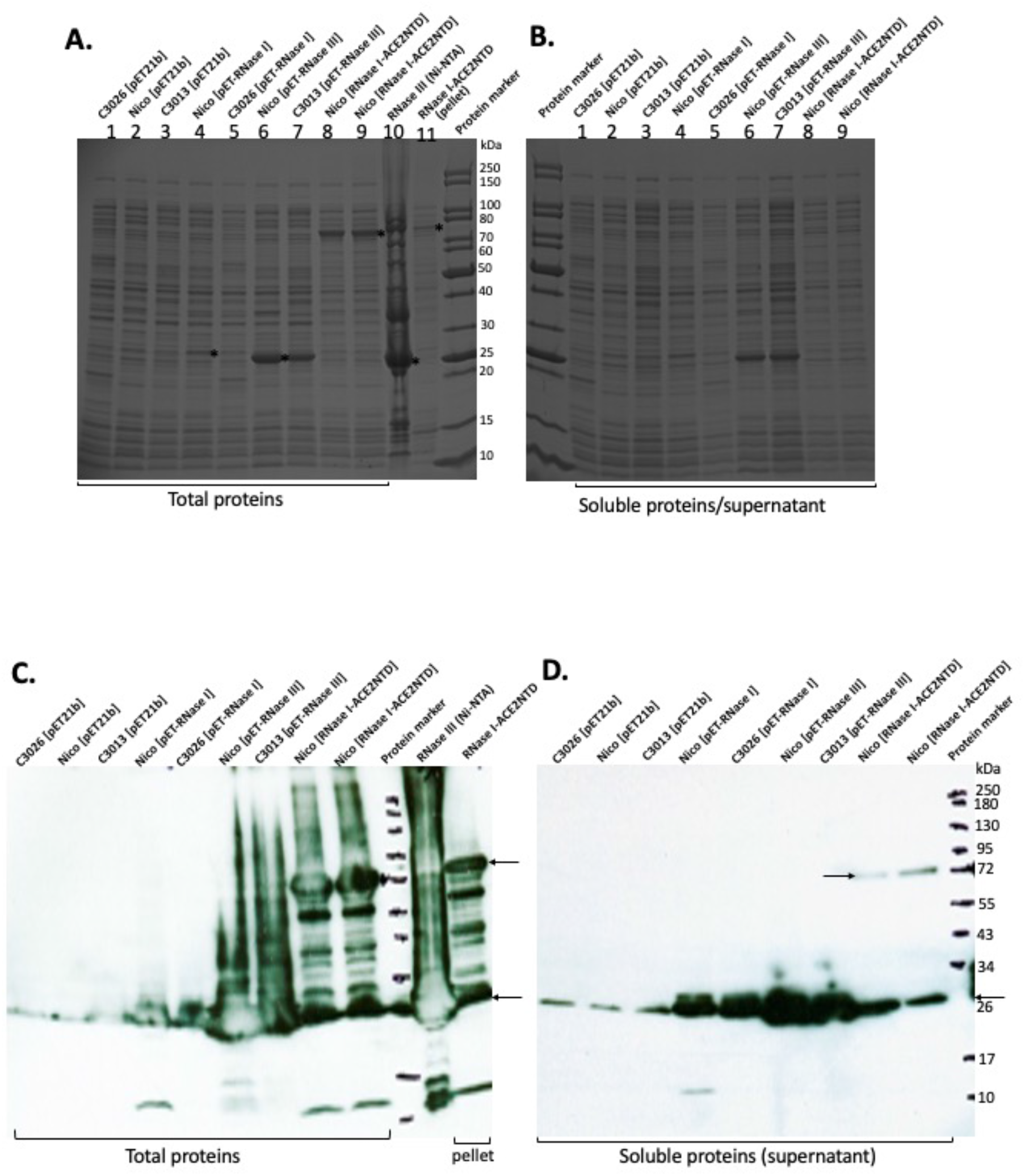
SDS-PAGE and Western blot analysis of *E. coli* lysate expressing RNase I (6xHis), RNase III (6xHis), and RNase I-ACE2NTD (6xHis) fusion. **A.** Total proteins in cell lysate. **B.** Soluble proteins in the supernatant after centrifugation in a microcentrifuge (10k rmp/min, 15 min). **C** and **D**. 6xHis-tagged proteins detected by anti-6xHis Ab in Western blot. Arrow and “*” indicate the target proteins. Color pre-stained protein marker (not shown) was from NEB.

## Reference

[1] R.R. Gaddam, S. Chambers, and M. Bhatia, ACE and ACE2 in inflammation: a tale of two enzymes. Inflamm Allergy Drug Targets 13 (2014) 224–34.

[2] R.A.S. Santos, G.Y. Oudit, T. Verano-Braga, G. Canta, U.M. Steckelings, and M. Bader, The renin-angiotensin system: going beyond the classical paradigms. Am J Physiol Heart Circ Physiol 316 (2019) H958–H970.

[3] W. Li, M.J. Moore, N. Vasilieva, J. Sui, S.K. Wong, M.A. Berne, M. Somasundaran, J.L. Sullivan, K. Luzuriaga, T.C. Greenough, H. Choe, and M. Farzan, Angiotensin-converting enzyme 2 is a functional receptor for the SARS coronavirus. Nature 426 (2003) 450–4.

[4] M. Hoffmann, H. Kleine-Weber, S. Schroeder, N. Kruger, T. Herrler, S. Erichsen, T.S. Schiergens, G. Herrler, N.H. Wu, A. Nitsche, M.A. Muller, C. Drosten, and S. Pohlmann, SARS-CoV-2 Cell Entry Depends on ACE2 and TMPRSS2 and Is Blocked by a Clinically Proven Protease Inhibitor. Cell (2020).

[5] C. Wang, P.W. Horby, F.G. Hayden, and G.F. Gao, A novel coronavirus outbreak of global health concern. Lancet 395 (2020) 470–473.

[6] Q. Li, X. Guan, P. Wu, X. Wang, L. Zhou, Y. Tong, R. Ren, K.S.M. Leung, E.H.Y. Lau, J.Y. Wong, X. Xing, N. Xiang, Y. Wu, C. Li, Q. Chen, D. Li, T. Liu, J. Zhao, M. Liu, W. Tu, C. Chen, L. Jin, R. Yang, Q. Wang, S. Zhou, R. Wang, H. Liu, Y. Luo, Y. Liu, G. Shao, H. Li, Z. Tao, Y. Yang, Z. Deng, B. Liu, Z. Ma, Y. Zhang, G. Shi, T.T.Y. Lam, J.T. Wu, G.F. Gao, B.J. Cowling, B. Yang, G.M. Leung, and Z. Feng, Early Transmission Dynamics in Wuhan, China, of Novel Coronavirus-Infected Pneumonia. N Engl J Med 382 (2020) 1199–1207.

[7] J. Paget, P. Spreeuwenberg, V. Charu, R.J. Taylor, A.D. Iuliano, J. Bresee, L. Simonsen, C. Viboud, N. Global Seasonal Influenza-associated Mortality Collaborator, and G.L.C. Teams*, Global mortality associated with seasonal influenza epidemics: New burden estimates and predictors from the GLaMOR Project. J Glob Health 9 (2019) 020421.

[8] M.A. Lake, What we know so far: COVID-19 current clinical knowledge and research. Clin Med (Lond) 20 (2020) 124–127.

[9] P. Mehta, D.F. McAuley, M. Brown, E. Sanchez, R.S. Tattersall, J.J. Manson, and U.K. Hlh Across Speciality Collaboration, COVID-19: consider cytokine storm syndromes and immunosuppression. Lancet 395 (2020) 1033–1034.

[10] C. Huang, Y. Wang, X. Li, L. Ren, J. Zhao, Y. Hu, L. Zhang, G. Fan, J. Xu, X. Gu, Z. Cheng, T. Yu, J. Xia, Y. Wei, W. Wu, X. Xie, W. Yin, H. Li, M. Liu, Y. Xiao, H. Gao, L. Guo, J. Xie, G. Wang, R. Jiang, Z. Gao, Q. Jin, J. Wang, and B. Cao, Clinical features of patients infected with 2019 novel coronavirus in Wuhan, China. Lancet 395 (2020) 497–506.

[11] V. Monteil, H. Kwon, P. Prado, A. Hagelkruys, R.A. Wimmer, M. Stahl, A. Leopoldi, E. Garreta, C. Hurtado Del Pozo, F. Prosper, J.P. Romero, G. Wirnsberger, H. Zhang, A.S. Slutsky, R. Conder, N. Montserrat, A. Mirazimi, and J.M. Penninger, Inhibition of SARS-CoV-2 Infections in Engineered Human Tissues Using Clinical-Grade Soluble Human ACE2. Cell 181 (2020) 905–913 e7.

[12] J.K. Millet, and G.R. Whittaker, Physiological and molecular triggers for SARS-CoV membrane fusion and entry into host cells. Virology 517 (2018) 3–8.

[13] S. Wang, F. Guo, K. Liu, H. Wang, S. Rao, P. Yang, and C. Jiang, Endocytosis of the receptor-binding domain of SARS-CoV spike protein together with virus receptor ACE2. Virus Res 136 (2008) 8–15.

[14] B. Tripet, M.W. Howard, M. Jobling, R.K. Holmes, K.V. Holmes, and R.S. Hodges, Structural characterization oi the SARS-coronavirus spike S fusion protein core. J Biol Chem 279 (2004) 20836–49.

[15] S.K. Wong, W. Li, M.J. Moore, H. Choe, and M. Farzan, A 193-amino acid fragment of the SARS coronavirus S protein efficiently binds angiotensin-converting enzyme 2. J Biol Chem 279 (2004) 3197–201.

[16] X. Ou, Y. Liu, X. Lei, P. Li, D. Mi, L. Ren, L. Guo, R. Guo, T. Chen, J. Hu, Z. Xiang, Z. Mu, X. Chen, J. Chen, K. Hu, Q. Jin, J. Wang, and Z. Qian, Characterization of spike glycoprotein of SARS-CoV-2 on virus entry and its immune cross-reactivity with SARS-CoV. Nature communications 11 (2020) 1620.

[17] L. Du, Y. He, Y. Zhou, S. Liu, B.J. Zheng, and S. Jiang, The spike protein of SARS-CoV--a target for vaccine and therapeutic development. Nat Rev Microbiol 7 (2009) 226–36.

[18] D. Wrapp, N. Wang, K.S. Corbett, J.A. Goldsmith, C.L. Hsieh, O. Abiona, B.S. Graham, and J.S. McLellan, Cryo-EM structure of the 2019-nCoV spike in the prefusion conformation. Science 367 (2020) 1260–1263.

[19] R. Yan, Y. Zhang, Y. Li, L. Xia, Y. Guo, and Q. Zhou, Structural basis for the recognition of SARS-CoV-2 by full-length human ACE2. Science 367 (2020) 1444–1448.

[20] S.A. Kemp, D.A. Collier, R. Datir, I. Ferreira, S. Gayed, A. Jahun, M. Hosmillo, C. Rees-Spear, P. Mlcochova, I.U. Lumb, D.J. Roberts, A. Chandra, N. Temperton, K. Sharrocks, E. Blane, J. Briggs, M.J. van Gils, K. Smith, J.R. Bradley, C. Smith, R. Doffinger, L. Ceron-Gutierrez, G. Barcenas-Morales, D.D. Pollock, R.A. Goldstein, A. Smielewska, J.P. Skittrall, T. Gouliouris, I.G. Goodfellow, E. Gkrania-Klotsas, C. Illingworth, L.E. McCoy, and R.K. Gupta, Neutralising antibodies in Spike mediated SARS-CoV-2 adaptation. medRxiv (2020).

[21] M. Hoffmann, H. Kleine-Weber, and S. Pohlmann, A Multibasic Cleavage Site in the Spike Protein of SARS-CoV-2 Is Essential for Infection of Human Lung Cells. Mol Cell 78 (2020) 779–784 e5.

[22] I. Glowacka, S. Bertram, M.A. Muller, P. Allen, E. Soilleux, S. Pfefferle, I. Steffen, T.S. Tsegaye, Y. He, K. Gnirss, D. Niemeyer, H. Schneider, C. Drosten, and S. Pohlmann, Evidence that TMPRSS2 activates the severe acute respiratory syndrome coronavirus spike protein for membrane fusion and reduces viral control by the humoral immune response. J Virol 85 (2011) 4122–34.

[23] S. Matsuyama, N. Nao, K. Shirato, M. Kawase, S. Saito, I. Takayama, N. Nagata, T. Sekizuka, H. Katoh, F. Kato, M. Sakata, M. Tahara, S. Kutsuna, N. Ohmagari, M. Kuroda, T. Suzuki, T. Kageyama, and M. Takeda, Enhanced isolation of SARS-CoV-2 by TMPRSS2-expressing cells. Proc Natl Acad Sci U S A 117 (2020) 7001–7003.

[24] K.H. Chan, J.S. Peiris, S.Y. Lam, L.L. Poon, K.Y. Yuen, and W.H. Seto, The Effects of Temperature and Relative Humidity on the Viability of the SARS Coronavirus. Adv Virol 2011 (2011) 734690.

[25] N. van Doremalen, T. Bushmaker, D.H. Morris, M.G. Holbrook, A. Gamble, B.N. Williamson, A. Tamin, J.L. Harcourt, N.J. Thornburg, S.I. Gerber, J.O. Lloyd-Smith, E. de Wit, and V.J. Munster, Aerosol and Surface Stability of SARS-CoV-2 as Compared with SARS-CoV-1. N Engl J Med (2020).

[26] S.F. Ahmed, A.A. Quadeer, and M.R. McKay, Preliminary Identification of Potential Vaccine Targets for the COVID-19 Coronavirus (SARS-CoV-2) Based on SARS-CoV Immunological Studies. Viruses 12 (2020).

[27] M.J. Mulligan, K.E. Lyke, N. Kitchin, J. Absalon, A. Gurtman, S. Lockhart, K. Neuzil, V. Raabe, R. Bailey, K.A. Swanson, P. Li, K. Koury, W. Kalina, D. Cooper, C. Fontes-Garfias, P.Y. Shi, O. Tureci, K.R. Tompkins, E.E. Walsh, R. Frenck, A.R. Falsey, P.R. Dormitzer, W.C. Gruber, U. Sahin, and K.U. Jansen, Phase I/II study of COVID-19 RNA vaccine BNT162b1 in adults. Nature 586 (2020) 589–593.

[28] E.E. Walsh, R.W. Frenck, Jr., A.R. Falsey, N. Kitchin, J. Absalon, A. Gurtman, S. Lockhart, K. Neuzil, M.J. Mulligan, R. Bailey, K.A. Swanson, P. Li, K. Koury, W. Kalina, D. Cooper, C. Fontes-Garfias, P.Y. Shi, O. Tureci, K.R. Tompkins, K.E. Lyke, V. Raabe, P.R. Dormitzer, K.U. Jansen, U. Sahin, and W.C. Gruber, Safety and Immunogenicity of Two RNA-Based Covid-19 Vaccine Candidates. N Engl J Med 383 (2020) 2439–2450.

[29] E.J. Anderson, N.G. Rouphael, A.T. Widge, L.A. Jackson, P.C. Roberts, M. Makhene, J.D. Chappell, M.R. Denison, L.J. Stevens, A.J. Pruijssers, A.B. McDermott, B. Flach, B.C. Lin, N.A. Doria-Rose, S. O’Dell, S.D. Schmidt, K.S. Corbett, P.A. Swanson, 2nd, M. Padilla, K.M. Neuzil, H. Bennett, B. Leav, M. Makowski, J. Albert, K. Cross, V.V. Edara, K. Floyd, M.S. Suthar, D.R. Martinez, R. Baric, W. Buchanan, C.J. Luke, V.K. Phadke, C.A. Rostad, J.E. Ledgerwood, B.S. Graham, J.H. Beigel, and R.N.A.S.G. m, Safety and Immunogenicity of SARS-CoV-2 mRNA-1273 Vaccine in Older Adults. N Engl J Med 383 (2020) 2427–2438.

[30] L.A. Jackson, E.J. Anderson, N.G. Rouphael, P.C. Roberts, M. Makhene, R.N. Coler, M.P. McCullough, J.D. Chappell, M.R. Denison, L.J. Stevens, A.J. Pruijssers, A. McDermott, B. Flach, N.A. Doria-Rose, K.S. Corbett, K.M. Morabito, S. O’Dell, S.D. Schmidt, P.A. Swanson, 2nd, M. Padilla, J.R. Mascola, K.M. Neuzil, H. Bennett, W. Sun, E. Peters, M. Makowski, J. Albert, K. Cross, W. Buchanan, R. Pikaart-Tautges, J.E. Ledgerwood, B.S. Graham, J.H. Beigel, and R.N.A.S.G. m, An mRNA Vaccine against SARS-CoV-2 - Preliminary Report. N Engl J Med 383 (2020) 1920–1931.

[31] L.A. Jackson, P.C. Roberts, and B.S. Graham, A SARS-CoV-2 mRNA Vaccine - Preliminary Report. Reply. N Engl J Med 383 (2020) 1191–1192.

[32] M.D. Knoll, and C. Wonodi, Oxford-AstraZeneca COVID-19 vaccine efficacy. Lancet 397 (2021) 72–74.

[33] D.E. Gordon, G.M. Jang, M. Bouhaddou, J. Xu, K. Obernier, K.M. White, M.J. O’Meara, V.V. Rezelj, J.Z. Guo, D.L. Swaney, T.A. Tummino, R. Huttenhain, R.M. Kaake, A.L. Richards, B. Tutuncuoglu, H. Foussard, J. Batra, K. Haas, M. Modak, M. Kim, P. Haas, B.J. Polacco, H. Braberg, J.M. Fabius, M. Eckhardt, M. Soucheray, M.J. Bennett, M. Cakir, M.J. McGregor, Q. Li, B. Meyer, F. Roesch, T. Vallet, A. Mac Kain, L. Miorin, E. Moreno, Z.Z.C. Naing, Y. Zhou, S. Peng, Y. Shi, Z. Zhang, W. Shen, I.T. Kirby, J.E. Melnyk, J.S. Chorba, K. Lou, S.A. Dai, I. Barrio-Hernandez, D. Memon, C. Hernandez-Armenta, J. Lyu, C.J.P. Mathy, T. Perica, K.B. Pilla, S.J. Ganesan, D.J. Saltzberg, R. Rakesh, X. Liu, S.B. Rosenthal, L. Calviello, S. Venkataramanan, J. Liboy-Lugo, Y. Lin, X.P. Huang, Y. Liu, S.A. Wankowicz, M. Bohn, M. Safari, F.S. Ugur, C. Koh, N.S. Savar, Q.D. Tran, D. Shengjuler, S.J. Fletcher, M.C. O’Neal, Y. Cai, J.C.J. Chang, D.J. Broadhurst, S. Klippsten, P.P. Sharp, N.A. Wenzell, D. Kuzuoglu-Ozturk, H.Y. Wang, R. Trenker, J.M. Young, D.A. Cavero, J. Hiatt, T.L. Roth, U. Rathore, A. Subramanian, J. Noack, M. Hubert, R.M. Stroud, A.D. Frankel, O.S. Rosenberg, K.A. Verba, D.A. Agard, M. Ott, M. Emerman, N. Jura, et al., A SARS-CoV-2 protein interaction map reveals targets for drug repurposing. Nature 583 (2020) 459–468.

[34] P. Riggs, Expression and purification of recombinant proteins by fusion to maltose-binding protein. Mol Biotechnol 15 (2000) 51–63.

[35] S. Nallamsetty, and D.S. Waugh, Solubility-enhancing proteins MBP and NusA play a passive role in the folding of their fusion partners. Protein Expr Purif 45 (2006) 175–82.

[36] S. Padmanabhan, K. Zhou, C.Y. Chu, R.W. Lim, and L.W. Lim, Overexpression, biophysical characterization, and crystallization of ribonuclease I from Escherichia coli, a broad-specificity enzyme in the RNase T2 family. Arch Biochem Biophys 390 (2001) 42–50.

[37] D.H. Bechhofer, and M.P. Deutscher, Bacterial ribonucleases and their roles in RNA metabolism. Crit Rev Biochem Mol Biol 54 (2019) 242–300.

[38] J. Xiao, C.E. Feehery, G. Tzertzinis, and C.V. Maina, E. coli RNase III(E38A) generates discrete-sized products from long dsRNA. RNA 15 (2009) 984–91.

[39] S. Sorrentino, M. Naddeo, A. Russo, and G. D’Alessio, Degradation of double-stranded RNA by human pancreatic ribonuclease: crucial role of noncatalytic basic amino acid residues. Biochemistry 42 (2003) 10182–90.

[40] P. Koczera, L. Martin, G. Marx, and T. Schuerholz, The Ribonuclease A Superfamily in Humans: Canonical RNases as the Buttress of Innate Immunity. Int J Mol Sci 17 (2016).

[41] R. Lu, X. Zhao, J. Li, P. Niu, B. Yang, H. Wu, W. Wang, H. Song, B. Huang, N. Zhu, Y. Bi, X. Ma, F. Zhan, L. Wang, T. Hu, H. Zhou, Z. Hu, W. Zhou, L. Zhao, J. Chen, Y. Meng, J. Wang, Y. Lin, J. Yuan, Z. Xie, J. Ma, W.J. Liu, D. Wang, W. Xu, E.C. Holmes, G.F. Gao, G. Wu, W. Chen, W. Shi, and W. Tan, Genomic characterisation and epidemiology of 2019 novel coronavirus: implications for virus origins and receptor binding. Lancet 395 (2020) 565–574.

[42] L. Lubbe, G.E. Cozier, D. Oosthuizen, K.R. Acharya, and E.D. Sturrock, ACE2 and ACE: structure-based insights into mechanism, regulation and receptor recognition by SARS-CoV. Clin Sci (Lond) 134 (2020) 2851–2871.

[43] Y. Alguel, J. Leung, S. Singh, R. Rana, L. Civiero, C. Alves, and B. Byrne, New tools for membrane protein research. Curr Protein Pept Sci 11 (2010) 156–65.

[44] R.T. Raines, Ribonuclease A. Chem Rev 98 (1998) 1045–1066.

[45] P. Klemm, and M.A. Schembri, Fimbriae-assisted bacterial surface display of heterologous peptides. Int J Med Microbiol 290 (2000) 215–21.

[46] K. Jacoby, M. Metzger, B.W. Shen, M.T. Certo, J. Jarjour, B.L. Stoddard, and A.M. Scharenberg, Expanding LAGLIDADG endonuclease scaffold diversity by rapidly surveying evolutionary sequence space. Nucleic acids research (2012).

[47] G.J. Gopal, and A. Kumar, Strategies for the production of recombinant protein in Escherichia coli. Protein J 32 (2013) 419–25.

[48] X. He, V. Hull, J.A. Thomas, X. Fu, S. Gidwani, Y.K. Gupta, L.W. Black, and S.Y. Xu, Expression and purification of a single-chain Type IV restriction enzyme Eco94GmrSD and determination of its substrate preference. Sci Rep 5 (2015) 9747.

